# AUTOMATIC EXTRACTION OF ACTIN NETWORKS IN PLANTS

**DOI:** 10.1101/2023.01.18.524528

**Authors:** Jordan Hembrow, Michael J. Deeks, David M. Richards

**Affiliations:** Living Systems Institute and Department of Physics and Astronomy, University of Exeter, Exeter EX4 4QD, UK; Biosciences, University of Exeter, Exeter, EX4 4PY

**Keywords:** Actin, Cytoskeleton, *Arabidopsis*, Image Analysis, Network Extraction, DRAGoN

## Abstract

The actin cytoskeleton is essential in eukaryotes, not least in the plant kingdom where it plays key roles in cell expansion, cell division, environmental responses and pathogen defence. Yet, the precise structure-function relationships of properties of the actin network in plants are still to be unravelled, including details of how the network configuration depends upon cell type, tissue type and developmental stage. Part of the problem lies in the difficulty of extracting high-quality, three-dimensional, quantitative measures of actin network features from microscopy data. To address this problem, we have developed DRAGoN, a novel image analysis algorithm that can automatically extract the actin network across a range of cell types, providing seventeen different quantitative measures that describe the network at a local level. Using this algorithm, we then studied a number of cases in *Arabidopsis thaliana*, including several different tissues, a variety of actin-affected mutants, and cells responding to powdery mildew. In many cases we found statistically-significant differences in actin network properties. In addition to these results, our algorithm is designed to be easily adaptable to other tissues, mutants and plants, and so will be a valuable asset for the study and future biological engineering of the actin cytoskeleton in globally-important crops.

## 1 Introduction

Being able to extract spatial networks composed of one-dimensional structures, from road networks to sub-cellular biological filaments, is a recurring theme throughout many research areas. Nowhere is this more important than for the cytoskeleton, where resolution-limited imaging data can make network extraction extremely difficult. In addition, this problem is by far the most acute for actin, the narrowest element of the cytoskeleton, which is ubiquitous throughout eukaryotic cells.

### 1.1 The Cytoskeleton

Amongst a multitude of other functions, the cytoskeletal network, prevalent in almost all eukaryotic cells, provides physical shape and structure to cells, aids in cell growth, and plays a key role in trafficking[1]. Formed from polymerisation of discrete protein sub-units, the cytoskeleton connects to various organelles (including the nucleus) and the plasma membrane. This modular nature of the cytoskeleton allows it to be dynamic, adapting as necessary to environmental changes relayed via a host of signalling processes[2]. Many of the cytoskeletal sub-units have been highly conserved during evolution and are found in most eukaryotic cells, with homologues even present in some prokaryotes[3].

The cytoskeleton is typically divided into three distinct components: microtubules, intermediate filaments and actin[4, 5]. Microtubules, the cytoskeletal component with the widest cross-section at about 25nm in diameter, are hollow tubes consisting of repeated *α*- and *β*-tubulin sub-units[6]. In animals and fungi, they play a number of roles including aiding in the formation of flagella or cilia[7], providing structures for material transport, and positioning of the mitotic spindle during cell division[8]. In plants, microtubules retain a role in cell division but also guide cell wall development through their relationship with wall-building enzyme complexes in the plasma membrane[9].

Unlike the globular units of microtubules and actin filaments, intermediate filaments are themselves constructed from filamentous sub-units, and confer strength as well as stress resistance to the cell[10]. Although they are present in almost all mammalian cells, their existence in plants is still hotly debated[11].

Finally, actin filaments, also known as microfilaments, are the narrowest components of the cytoskeleton and are constructed from globular actin sub-units (G-actin) that assemble to form a helical structure 5-7nm in diameter[12]. In combination with myosin motors, actin aids in transport by providing the roads and pathways for cellular cargo[13]. Actin microfilaments are present as both individual filaments and bundled into thicker filaments, and play a key role in plant cell growth and internal transport[14]. It is actin that will be our primary concern here, although the approach we develop is likely to be easily adaptable to other cytoskeletal components.

### 1.2 Actin in Plants

Although actin is present across eukaryotes, there are important differences between kingdoms, not least in plants. The versatility of actin enables it to play a key role in many aspects of plant life, including root tip growth[15] and cell division[16]. The ubiquity of the actin cytoskeleton, alongside the fact it can be constantly remodelled in response to external forces and environmental factors, means that active, directional transport is always available throughout plant cells. This transport is particularly fast in plants, with Golgi speeds reaching up to 7μm s^−1^ [17]. To enable these speeds the actin cytoskeleton typically runs mainly parallel to the long axis of an expanded cell, sandwiched between the vacuole and the plasma membrane and perpendicular to the orientation of microtubules[18]. The network consists of two distinct populations: the fine F-actin arrays that undergo rapid and stochastic remodelling with polymerisation rates in *A. thaliana* hypocotyls reaching 1.7μm s^−1^ [19], and the thicker bundles responsible for long-range transport.

The actin cytoskeleton in plants is regulated through a large array of genes and proteins that are coordinated by numerous signalling pathways. External influences on the cell, whether from environmental factors (such as light or temperature) or other organisms (such as pathogens like *Blumeria graminis* that can attack the cell well through enzymes or physical forces), can stimulate these pathways, triggering adaptations in the actin network to counter the stimulus. Actin plays a key role in the response to pathogen attack, rapidly remodelling itself at the site of the plasma membrane where the pathogen makes contact[20, 21]. F-actin aggregation at the attempted penetration site, which enables cytoplasm accumulation[22], can occur within approximately 20 seconds and can be triggered by small applied forces of only 4μN[23]). Depolymerisation of actin using drug treatments causes the catastrophic failure of this focused penetration defence[24]. Conversely, crop breeders have taken advantage of a mutation named *mlo* that enhances the actin response in barley to offer increased protection against some types of fungi[25]. Pathogens are under selective pressure to adapt their repertoire of secreted effector proteins to suppress plant defences, including the actin cytoskeleton[26]. Climate change is forcing pathogens to migrate globally and is leading to novel encounters between agricultural cultivars of plant and virulent strains of microbes[27], which adds urgency to the need to understand how plant defences can be enhanced. The ability to detect and measure these changes in actin structure and dynamics will be critical in understanding the defensive systems that plants deploy.

### 1.3 Properties of Actin

Monomers of actin (G-actin) can be assembled into filamentous actin polymers (F-actin), initially through the nucleation of G-actin into a trimer, before additional G-actin units are sequentially added (see Fig. 1)[28]. One end of the filament (called the barbed or positive end) has an exposed ATP binding site that is missing from the other end (the pointed or negative end), thus giving the structure an overall polarity[29]. While polymerisation occurs at both ends, the barbed end has a significantly increased rate of assembly. G-actin monomers can also leave both ends, resulting in disassembly. G-actin monomers bound to ATP are more likely to be assembled, shortly after which the ATP is converted to ADP-P_i_ through nucleotide hydrolysis. After a delay, estimated to be around the order of seconds[30], the P_i_ is then released, leaving just ADP in the binding site[31]. The various ADP-ATP combinations contained within the nucleated monomers gives an indication of the age of that part of the filament, which can be detected by disassembly machinery such as Actin Depolymerising Factors (ADFs; also known as cofilins)[32]. Filament polarity results in frequent growth at the barbed end and disassembly at the pointed end, known as treadmilling, which allows the structure to be adaptive to changes in conditions and external forces. F-actin can also bundle together, forming thicker and straighter filamentous structures that are more stable and have a longer lifetime. These cross-linking and bundling processes are initiated by proteins such as villin and fimbrin[33].

**Figure 1:**
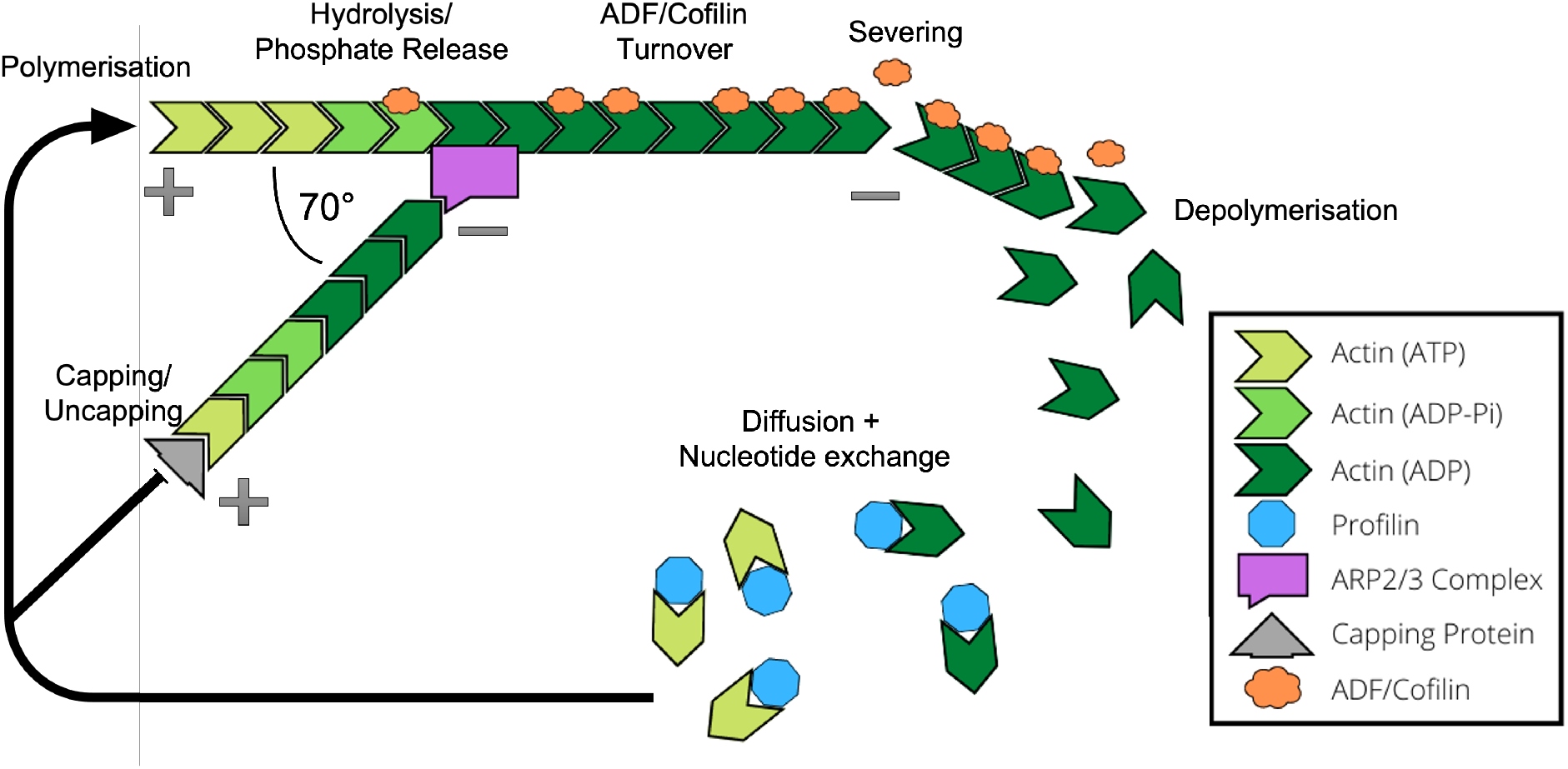
Actin network dynamics. Actin within the cell consists of both single monomers (G-actin) and branched filaments (F-actin). Filaments have an orientation, with a pointed end (-) and a barbed end (+). Polymerisation and disassembly can occur at both ends, but polymerisation typically occurs at a higher rate at the barbed end, while disassembly is more common at the pointed end. A range of proteins and signalling pathways contribute to the turnover of actin monomers, which drives the structure of the actin cytoskeleton and enables a high level of organisation and adaptation. The Arp2/3 complex is involved in filament nucleation and branching, while capping proteins bind to the barbed end to prevent both polymerisation and depolymerisation. Severing proteins such as ADF/cofilin can sever the filament creating an additional barbed and pointed end.

A range of actin binding proteins (ABPs) work in harmony with the cytoskeleton to enable a higher level of organisation (see Fig. 1). For example, the actin related protein (ARP) 2/3 complex is an actin nucleator[34, 35], which works alongside formin proteins and other nucleators to aid in the control of the rate and timing of actin assembly and branching[36, 37]. ADF/cofilin and cyclase associated protein (CAP) bind to G-actin and help to regulate the available pool of actin monomers for assembly[38]. Capping proteins bind to the barbed end of F-actin and prevent both assembly and disassembly, thus controlling length and growth. Conversely, severing proteins such as ADF/cofilin and villin sever filaments[39], usually towards the pointed end where the actin is bound to ADP[40] and so shorten the filament whilst creating a new barbed and pointed end.

Profilins are proteins that work in collaboration with plant formins to regulate several aspects of cytoskeletal development and growth. *A. thaliana* contains five isoforms of profilin, of which AtPRF3 has an atypical N-terminal extension that enhances the affinity for the polyproline region of formin AtFH1. AtPRF3 binding to formin and subsequent oligermerisation has been shown to inhibit AtFH1-mediated actin nucleation. In response to pathogen associated molecular patterns, the transcription of AtPRF3 and protein degradation are altered in order to modulate actin turnover[41]. Furthermore, formin AtFH4/FORMIN4 has been found to interact with AtPRF2[42]. During plant immune responses to fungi and oomycetes this contributes to cytoskeletal remodelling and ensures the maintenance of the filamentous actin arrays in proximity to sites of attempted penetration site[43, 44].

The numerous types of protein working in concert to keep the cytoskeletal network organised allows it to be adaptive and efficiently restructured as required. ADF[32], profilin[41], formin[44] and the ARP2/3 complex[43] have all been implicated in the changes in structure and dynamics that place the actin cytoskeleton in a defensive poise against microbes. Each will have a subtle input into the behaviour of the actin array and decoding these contributions provides information about the key control points driving defence.

### 1.4 Analysing the Actin Network Structure

Being able to observe and measure quantitatively the structure of the cytoskeleton is important for understanding wild-type behaviour, mutant phenotypes, the effect of drug treatments, and the cellular response to external stimuli such as pathogen attack. The fact that the cytoskeleton is highly adaptive and versatile, rapidly responding to changes when required, only makes this task more difficult, not least because some small perturbations (such as HopW1 from *Pseudomonas syringe*[45]) are able to cripple the entire network and, by extension, the cell itself.

Typically, progress in this area requires significant amounts of data. This has become increasingly viable due to the development of genetically encoded probes (e.g. GFP-Lifeact and GFP-Fimbrin) which can be used across a range of imaging modalities to give significant insight into the actin network structure and its turnover. Analysing such large datasets by hand is undesirable for at least three reasons: subconscious bias can skew results, human error is hard to avoid, and the time required can be prohibitive. What is needed instead is a consistent, automatic, reproducible, objective method that can quickly and quantitatively analyse network changes.

The constant remodelling and treadmilling of the actin cytoskeleton poses a problem for any analysis method. Live-cell imaging can be limited by exposure times that lead to motion-blur of highly dynamic features. Cell fixation methods can alleviate this, but only give a single snapshot in time and are prone to artefacts from the fixation process. Modalities ideal for capturing highly dynamic actin networks such as Total Internal Reflection (TIRF) microscopy can be limited in their ability to capture the complete three-dimensional network. This can be better achieved by forms of confocal microscopy, but here too there can be limitations such as poor Z resolution. No imaging system will be able to capture perfectly every detail of the actin network, which makes maximising the information available from current methods a priority. The algorithm we develop here is a step towards achieving this.

### 1.5 Previous Work in this Direction

The importance of the cytoskeleton across many areas of cell biology has resulted in significant interest in automatic methods that enable visualisation and extraction of the network. Here we review those that are most relevant to our work here. For a recent review that summarises many of these tools and the different methods they employ, see Özdemir and Reski[46].

Breuer et al. have developed an actin network extraction tool called CytoSeg[47, 48], which leverages a Python backend with GUI implementation in ImageJ. The actin network is described by a weighted graph (*i.e*. a mathematical network of nodes connected by weighted edges) which builds upon the DeFiNe method[49]. This yields powerful and simple analysis for transport networks, including routing and distances travelled, well-suited for analysis of vesicle and organelle transport. However, one limitation of this approach is that filaments are represented by straight lines, a simplification that can miss key properties of actin networks such as filament curvature. Not knowing the full network shape (with realistic non-straight filaments) can make network analysis more difficult and can miss subtle network changes that are often seen in mutants and drug treatments.

One of the most common methods utilised for cytoskeletal extraction is based on open active contour models[50, 51, 52]. These examine changes in the intensity gradient of an image in order to find ridges, which form the backbone of actin filaments. These backbones (often called “snakes”) are defined in terms of the energy of their contour, which is minimised in the gradient field in order to generate the filament segmentation. This process can be computationally expensive depending on the number of iterations required to achieve accuracy, but the results tend to be impressive in terms of accuracy (precision and recall)[53]. Several tools are available which use these models, including imageJ plugins like JFilament[54] and standalone programs such as SOAX[55] and TSOAX[53]. Depending on the implementation, one drawback of these methods is that they often require significant human input to aid in the labelling of filaments and discern crossings. This has to be done separately for each image, which diminishes the utility of automatic methods. The more automated methods, such as SOAX, provide the option for batch processing and therefore require reduced input from start to finish. The number and sensitivity of the input parameters, however, make it difficult to achieve consistently accurate results across a variety of images, cell types and scenarios. While the input parameters can be adjusted on a per-image bases, this may interfere with the validity of any comparisons and could introduce bias into the results.

A complementary approach, based on the molecular mechanisms that govern actin network formation and dynamics, was developed by Schaub et al., and involves simulating the network itself and comparing the output to experimental images to determine the network properties[56]. Comparison between simulation results and experiments required knowledge of the imaging system point spread function, the resolution of the charge-coupled device (CCD), any non-uniform responses from pixels, and the magnification and numerical aperture (NA) of the lens. These methods were able give a good estimate of the underlying network, but only under the assumption that the simulation correctly captures the true underlying network dynamics. At present, with current knowledge of actin biophysics, this may not be the case. Further, fitting a many-parameter simulation to experimental data can both be time consuming and experience problems of model identifiability (where significantly different parameter values lead to similar network structures).

Several approaches have been developed for other imaging modalities, as well as software designed to identify other structures linked to the cytoskeleton. CellArchitect[57] is a tool designed to extract automatically plant microtubule data from large quantities of confocal microscopy data using Gaussian filters and local thresholding to segment the network from the background. From here, two-dimensional measures such as length, density and width can be extracted. SIFNE[58] is a tool designed to extract simple quantitative data (such as filament length and density) from the microtubule array imaged with single-molecule-localisation microscopy. ANNA-PALM[59] is another super-resolution-based tool that can reconstruct microtubule arrays, either via PALM or DNA-PAINT microscopy[60]. BundleTrac[61] and Actin Segmentation[62] both extract the network of actin filaments from Cryo-Electron Tomography images. BundleTrac requires manual input of “seed points” to start filament identificiation while Actin Segmentation is fully automated with analysis of the resulting networks yielding measurements of filament lengths, orientations and densities. Finally, FIESTA[63] and TipTracker[64] use epifluorescence microscopy to locate microtubules, with the former being shown to also work with TIRF microscopy (as does MTrack[65]).

Segmentation and analysis can be extended to additional biopolymers such as the endoplasmic reticulum. The MATLAB software AnalyzER[66] takes fluorescent labelled ER images from a confocal microscope (up to multi-channel 4D time-series data), enhances the tubular structures and then performs segmentation and skeletonisation. The skeleton is converted to a graph representation with nodes at junctions (containing data such as degree and branch angles) connected by edges (which also described length and average width). This approach was able to quantify the effects on the morphology of the ER under different drug treatments and abiotic stresses.

Depending on the assay, biopolymer of interest and imaging requirements, there are a range of techniques tested and available[67, 68, 69, 70, 71, 72, 73]. However, most of these solutions are tailored to specific forms of biological network and research questions, consequently they do not integrate a broad categorisation or quantification of individual network properties.

### 1.6 Plan of this paper

This paper is organised as follows. We first describe how the DRAGoN (<u>D</u>igital <u>R</u>econstruction of <u>A</u>ctin <u>G</u>l<u>o</u>bal <u>N</u>etworks) algorithm was developed and explain the seventeen quantitative measures that are then provided, with further detail provided in the Supporting Information. We also explain the various validation steps we performed to test and measure algorithm performance.

Next, we apply our algorithm to new data that we have collected from *A. thaliana*. This includes images of actin in wild-type (Col-0) cells, along with our *arp2-1* and *formin4/7/8* mutants, both in hypocotyls and in leaves. Further, we consider the effect of powdery mildew infection, which produces a localised response within infected cells.

Finally, we discuss the implications of the significant results that we found, both in the context of understanding the cytoskeleton and future research into plant immunity and food security. We conclude by discussing possible extensions to the algorithm as well as additional routes that could be taken for parameter tuning for various cell types and imaging modalities.

## 2 Results

### 2.1 Algorithm Development

Using an initial set of 20 images of GFP-Lifeact[74] labelled actin in wild-type *A. thaliana* (obtained as described in the Methods), we first developed the DRAGoN algorithm (available at https://github.com/JordanHembrow5/DRAGoN) for automatically extracting the actin skeleton from either a single 2D image or a 3D stack of images. We decided to base this in MATLAB due to the power of the image processing toolbox. MATLAB also has excellent backwards as well as forwards compatibility, ensuring easy adoption both by users with older versions and by future users with as yet unreleased versions.

Our algorithm follows five sequential steps (see Methods and the Supporting Information for full details): initial preparation, skeletonisation, image rotation, skeleton labelling and property measuring. We discuss the first four steps here, leaving the last for the next subsection.

For the first image preparation step, the images were filtered to reduce noise and eliminate uneven background illumination. Any filtering process will result in the loss of information, but is necessary to facilitate extraction of the true actin signal. A range of filters and algorithms were tested, each with a range of parameters, in order to determine the most suitable option. Best results were found by using a top-hat filter with a spherical structuring element. A top hat filter subtracts the morphological opening (an erosion followed by a dilation) of the image from the original image. This is similar to the rolling ball background filter in ImageJ, a commonly-used approach to analyse biological images. This aids in noise removal as well as correcting non-uniform illumination. A radius much larger than the width of the thickest filament was found to be suitable (between 2 to 3 times the width), otherwise filament signal may be removed or contrast at the edges may be lost. A Gaussian filter was also tested, but proved less successful as the filament edges were not preserved, making subsequent enhancements of tubular structures more difficult and measurements of thickness unreliable.

The second step was to convert the filtered image to a binary image of the skeletonised network, attempting to keep the filaments without any of the background. Simply applying a threshold to the raw image led to poor results (either struggling with detecting finer filaments or failing to adequately remove noise), so we first enhanced tubular structures using the MATLAB function *fibermetric* as shown in Fig. 2B. This works by calculating the Hessian of the image, determining its eigenvalues and searching for pixels with a small intensity change in one direction and a large change in the other[75]. Pixels satisfying this signify a tubular structure and so have their signal boosted. The benefit of this function is that it automatically adapts to various filament sizes, removing the need for any additional parameter tuning.

With the desired structures enhanced, we then created a binary image by using a threshold, shown in Fig. 2C. We tested both an adaptive and fixed threshold. For the adaptive threshold we used both the MATLAB default neighbourhood of around eighth of the image size as well as smaller sizes. However, this failed to perform any better than a fixed threshold, while being significantly more computationally expensive and requiring an additional parameter. It is possible that by fine tuning the neighbourhood size and the sensitivity, an adaptive threshold will produce an improved binary for some data sets, but this would require specific tuning for each image, something that would compromise comparisons between images. We therefore decided to use a fixed threshold, with the threshold value determined as a specific percentile of the image intensity distribution. We believe this approach reduces the sensitivity of the threshold parameter, enabling it to work better across multiple images and data sets, although it is worth noting that it is still the most sensitive parameter of our algorithm. A threshold intensity of between the 87^th^ and 92^nd^ percentile of the resultant image worked well in various scenarios, so we settled on 90% for the parameter value from this point forward.

**Figure 2:**
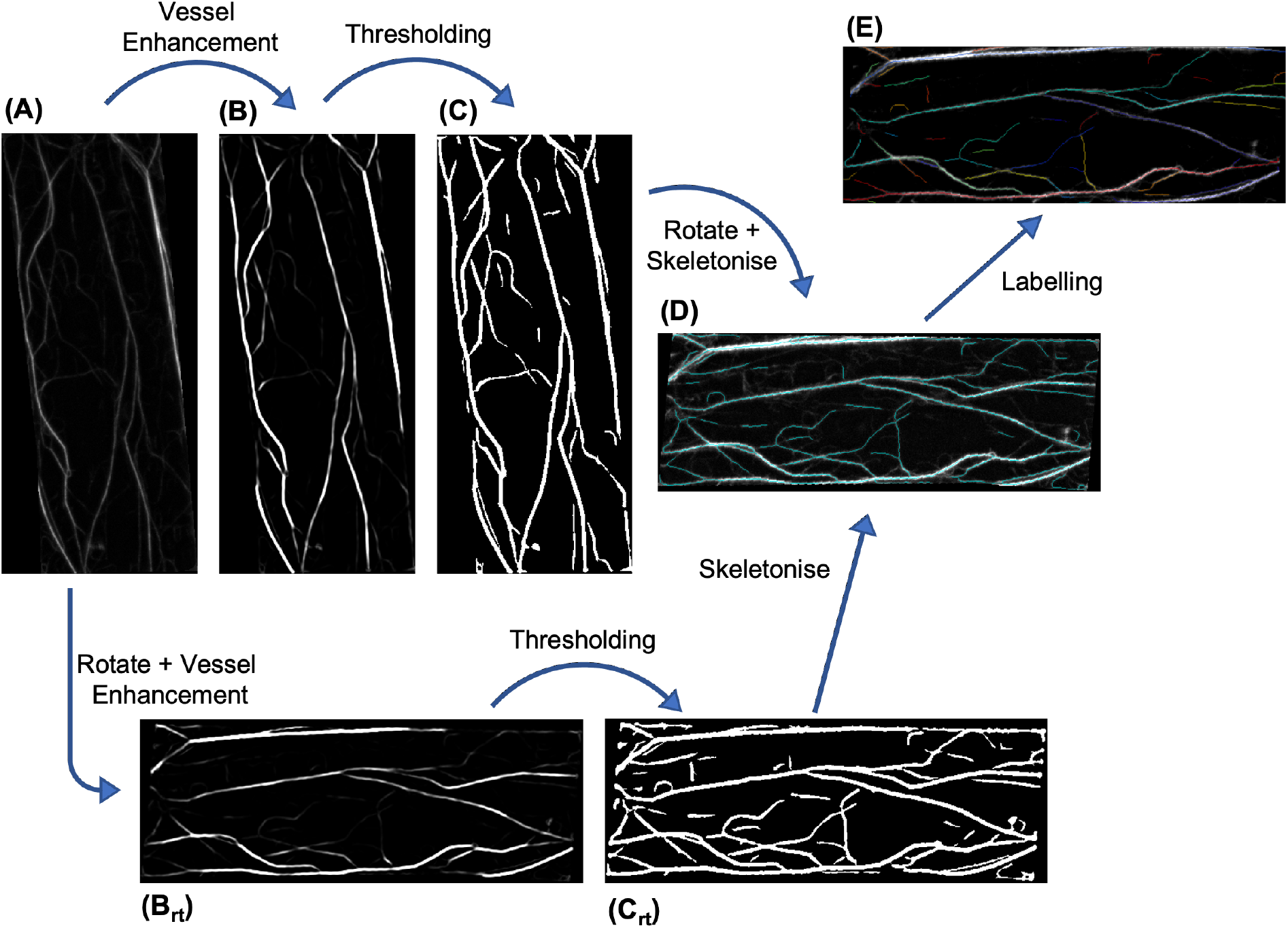
Skeletonisation Algorithm. Two parallel pathways are taken from the initial image (A) after background removal and filtering. The image had vessels enhanced (B) before thresholding into a binary image (C) before being skeletonised and rotated (D). Alongside this, the initial image was separately rotated before vessel enhancement (B_rt_) and thresholding (C_rt_) followed by skeletonisation (D). The skeleton results were combined to ensure no loss of information occurred in the rotation steps. From the combined skeleton, the labelling and relabelling processes were applied (E) to identify each filament and branch for further analysis

Despite filtering and parameter tuning there was still some remaining noise in the binary image. It was possible to remove some of this, before skeletonisation, by deleting objects with an area below some critical value. This critical value strongly depends on the imaging conditions and modality, and so would likely need to be determined independently for each given dataset.

The final part of the second step was the skeletonisation itself. To do this, pixels were removed from the perimeter of all binary objects until removing more would have altered the topology or Euler number (the number of objects in an image minus the number of holes). This yielded a single-pixel-thick line that represents the backbone of the actin network.

The third step involved image rotation. To simplify downstream analysis, images were rotated such that the long axis of the cell (or ROI) was parallel to the horizontal axis of the image. The rotation angle was determined by fitting an ellipse around the image mask and calculating the orientation of its major axis. When rotating discreet pixels, some positional accuracy is inevitably lost. If the rotation is performed before the binary stage, then this can be remedied by using a smoothing algorithm. However, if performed after skeletonisation then, as with a Gaussian filter, this would smooth the edges we have tried to preserve. The best solution we found to this was to implement the rotation step both before and after skeletonisation, and then combine the two outputs via a pixel-wise OR, as shown by the parallel workflows of Fig. 2. If after this step any filaments were thicker than one pixel, they were skeletonised again.

The fourth step was to label the skeleton so that individual filaments were identified by distinct positive integers and is shown in Fig. 2E. This required significant testing and development in order to eliminate edge cases. The details of each step taken to achieve this are described in detail in the Supporting Information. In brief, to leverage the flood-fill labelling algorithms in MATLAB, the filaments first had to be broken apart at every branch point so that branching filaments were given a different label ID to the main filament. It was found to be easier to also fragment the main filament in the process, and then rebuild and relabel afterwards. This was achieved by determining the labels around all (now removed) branch points, looking at the angles relative to each other, and rejoining the two that formed the straightest line.

### 2.2 Quantitative Measures

Once the actin skeleton had been extracted and correctly labelled, the fifth and final step involved extracting quantitative measures that described key properties of the network. It is these measures that can be used to characterise the actin cytoskeleton, providing a way of (i) understanding the underlying nature of the network and (ii) comparing different image sets (e.g. across cell types, mutants or drug treatments) in order to determine if there are significant differences. Seventeen types of measure are defined for 3D data and these are summarised in Table 1 and Fig. 3, as well as being described in detail in the Supporting Information. These measures are split into five different categories: whole cell properties, overall network properties, individual filament properties, curvature properties and branching properties, each of which is described below. Additional measures can easily be added if required at a later date.

**Table 1:** Quantitative network measures. The seventeen quantitative measures that we calculate for each extracted network along with a description of each. Full details are given in the Supporting Information. N=single number, L=list of numbers.

| Category | Name | Type | Code | Description |
| --- | --- | --- | --- | --- |
| Whole cell properties | Cell size | N | cellSize | Total number of pixels in the region of interest (ROI) |
|  | Cell orientation | N | orientation | Angle in radians that the image is rotated by such that the long axis of the cell is parallel to the x axis |
| Overall network properties | Skeleton density | N | skelDensity | Number of pixels in the skeleton divided by cell size |
|  | Structure size | L | Structures | Number of pixels in structures |
|  | Branch ratio | N | branchRatio | The number of branch points divided by the number of filaments |
|  | Branch point density | N | cellBPDensity | Number of branch points in the ROI divided by cell size |
| Individual filament properties | Filament width | L | filWidth | Average width of each filament |
|  | Filament lengths in 2D | L | filLenXY | Length of each filament in 2D, accounting for angles and curvature |
|  | Filament lengths in 3D | L | filLenXYZ | Length of each filament in 3D (the 2D length divided by the cosine of the filament angle relative to the imaging plane) |
|  | Filament angles in 2D | L | filamentAng.angXY | Angle of each filament relative to the x-axis |
|  | Filament angles in 3D | L | filamentAng.angZ | Angle of each filament relative to the confocal imaging plane; roughly estimated from endpoints and limited z-resolution |
|  | Average filament length | N | avgLen | Mean of the 3D filament lengths |
| Curvature properties | Signed filament curvature | L | curvatureSigned | Average signed curvature of each filament, measured over length scale $\lambda$ |
| | Unsigned filament curvature | L | curvature | Average unsigned curvature of each filament, measured over length scale $\lambda$ |
|  | Deviation | L | deviation | Average distance of each pixel in a filament from a straight line joining the endpoints |
| Branch properties | Branch angles in 2D | L | branchAng.angXY | Angle between main filament and a branch in 2D |
|  | Branch angles in 3D | L | branchAng.angZ | Angle between main filament and a branch in 3D |

**Figure 3:**
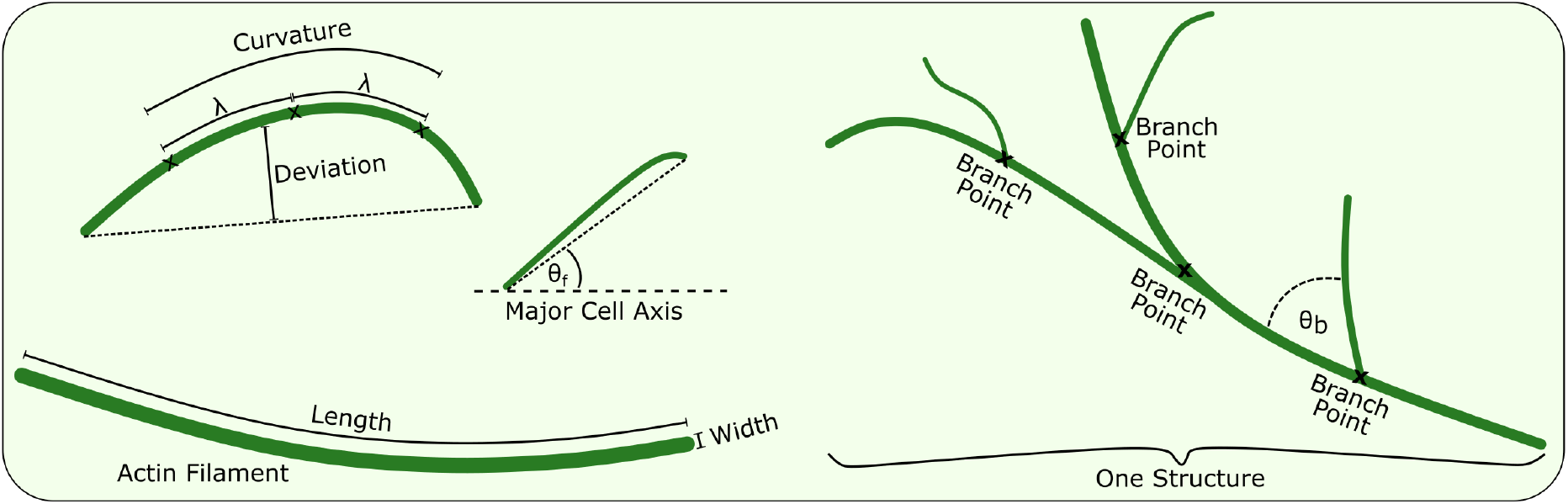
Network quantification. A visualisation of several of the measurements we define to quantify the actin network. The *cell size* is measured as the area encompassed by the black line of the outer bounding box and is used in the calculation of densities such as the skeleton and branch point density. The cell *orientation* is measured as the angle of the bounding box relative to the horizontal axis. The curvature, both the *signed filament curvature* and the *unsigned filament curvature*, are averaged over the whole length of each filament and measured via a series of three points separated by a characteristic length scale, λ. The *deviation* measures the average distance between a filament and a straight line connecting the end points. The *filament width* is defined for each filament as an average over the filament length. The *filament angle, θ_f_*, is measured relative to the major cell axis and is calculated both as a 2D and 3D version. The *branch angle*, *θ_b_*, is measured at every branch point and measures the deviation of the branch from the main filament; again, there is a 2D and 3D version. Finally, the *branch ratio* is the number of branch points divided by the number of labelled filaments.

#### Whole cell properties

The *cell size* is measured by counting the number of pixels in the ROI mask. This can then be used for density calculations in order to give quantities that are independent of the cell size. The *cell orientation* relative to the horizontal axis is measured by fitting the ROI to an ellipse and measuring the angle the ellipse makes to the horizontal axis. This measurement can then be used to align all cells to the horizontal axis to make further measurements more easily comparable.

#### Overall network properties

The *skeleton density* is measured as the total skeleton length in pixels divided by the cell area and can be used to highlight how tightly packed the network is. While many of the networks in our data sets were typically connected to each other, new disconnected filaments and structures could form. To measure this, we calculate the number of these structures and the mean *structure size*. A significant change in this value may indicate, for example, disruption to severing activity. Network branching frequency is measured using the *branch ratio*, calculated as the average number of branches per labelled filament, while the *branch point density* is calculated as the total number of branch points divided by the ROI area.

#### Individual filament properties

We also calculate a range of filament descriptors, including orientation, length and width. *Filament width* is defined as the average width in pixels and provides information about bundling activity. The *filament lengths in 2D* are measured by counting the number of pixels in the filament and adding 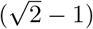 for every diagonal connection, then converting from pixels to physical length. For *filament lengths in 3D* these lengths are then scaled by cos *θ_z_* where *θ_z_* is the angle of displacement of the filament from the imaging plane. Changes in filament lengths may be indicative of growth, depolymerisation, or severing activity. The *filament angles in 2D* are obtained by calculating the angle relative to the horizontal axis of the line connecting the two filament end points. For *filament angles in 3D* the angle relative to the imaging plane is measured. The z-position of the filament end points is estimated by determining the position in the stack that contains the brightest pixel, which then allows the true angle to be measured. Finally, the *average filament length* is calculated as the mean of the 3D filament lengths.

#### Curvature properties

The curvature and non-linearity (deviation) of filaments are measured and represented by three different values. The curvature is measured over a fixed length scale λ in order to make the results independent of pixel size. For our data we chose λ = 475nm, which is equivalent to five pixels. This is large enough that individual noise at the pixel level does not adversely affect the results, and small enough that the calculated value is a good estimate of the local curvature. Further, this value should work for a range of common imaging modalities and resolutions. For all non-end points in a filament, a point either side is chosen as close to λ away as possible. These three points then have their Menger curvature calculated. The *signed filament curvature* is the absolute value of the average of these curvatures (including their signs). The *unsigned filament curvature* is the mean of the magnitudes of the curvatures (ignoring their signs). The non-linearity of the filament, referred to as the *deviation* from linear, was measured by calculating the absolute value of the mean distance between the filament and a straight line connecting the end points.

#### Branching properties

The *branch angles in 2D* are measured as the angle that a myosin motor would have to deviate to take the branch path instead of remaining on the main filament. This measurement is not calculated in the case that no branches are available. Finally, the *branch angles in 3D* is similar to the 2D measurement, but with the additional information of the z-position of the end points determined from the image stack. The distances between each pair of points are then calculated and used to determine the branch angle in 3D.

### 2.3 Algorithm Testing and Validation

Although testing the output of our algorithm against manual network segmentation gives some confidence that the algorithm can correctly extract the underlying actin network, it of course suffers from the problem that the human extraction may itself be faulty or biased. Rectifying this requires knowing the ground truth for a given network. To address this, we generated over 1,300 200×200px artificial images where the locations and shapes of all filaments were known perfectly in advance. Individual filaments were generated by drawing a smoothing spline between two randomly generated points and their midpoint, which was shifted from the middle by a random amount to ensure a curve was formed. In order to accurately represent various filament thicknesses, each image dimension was scaled by a factor of 19, reducing pixel size from 95nm^2^ (the pixel size in our microscopy assay) to 5nm^2^ such that F-actin can be represented by a pixel-thick line. A Gaussian blur was then applied to match the diffraction limit of our of microscopy assay, before the image was downscaled back to 200×200 pixels by taking the mean of each 19×19 block of pixels. To adjust the signal-to-noise ratio (SNR), the filament brightness was scaled to a desired value, then salt and pepper noise added and randomly (via a uniform distribution) scaled up to a maximum parameter value depending on the SNR level that was targeted. Finally, a Gaussian blur matching the diffraction limit was applied to the noise before the image of the filaments and noise were combined, yielding a simulated microscopy image. Several parameters throughout these steps were varied in order to mimic differences in image quality. This allowed us to test the robustness of the network extraction process. For more detail of this process, see the Supporting Information.

In order to quantify algorithm performance, two metrics were used: sensitivity and precision. These are calculated using the frequencies of true positives (TP), false positives (FP) and false negatives (FN). Sensitivity, also known as the true positive rate (TPR), describes the proportion of the network that is correctly detected and is given by TPR = TP/(TP + FN). Precision, also known as the positive predictive value (PPV), measures the positional accuracy of the identified network and is given by PPV = TP/ (TP + FP). As expected, there is a trade off between sensitivity and precision, which can be adjusted by altering various parameters, particularly the threshold parameter. By manually segmenting the real data shown in Fig. 2, we found a precision of 96% and a sensitivity of 72%, although these values are only indicative due to the difficulties and subjective nature of human segmentation.

First, by adjusting the maximum noise and filament brightness values, the signal-to-noise ratio (SNR) was varied. Because there is randomness associated with the image generation process, a range of SNR values correspond to a parameter set, and scaling between the parameters and the SNR is non-linear. Three different noise levels were generated (each with 100 images), each containing 10 filaments (see Fig. 4A). Precision remained at or above 97% across the three SNR levels, while the sensitivity drops from 89% at an SNR average of 7.3 to 86% at an average SNR of 4.3. For the low SNR group the sensitivity drops significantly to 57% for an average SNR of 3.4, highlighting the point of limitation for the algorithm and the data. Although our algorithm was designed to attempt to try to keep both precision and sensitivity as high as possible, it is possible to optimise for one of these by adjusting the binary threshold and/or the minimum filament size as explained below.

**Figure 4:**
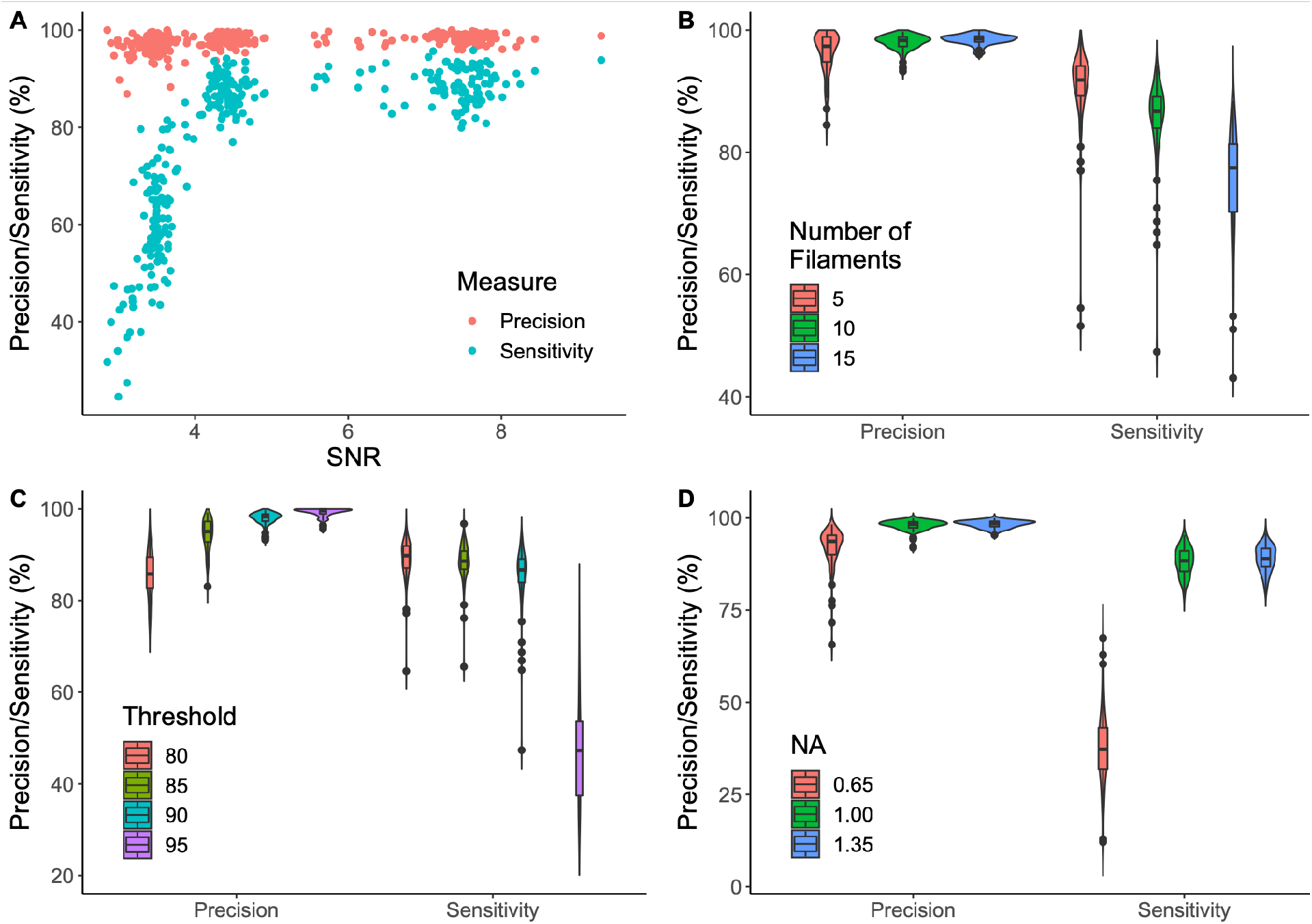
Testing network extraction performance. Artificially-generated data was analysed with the DRAGoN algorithm and the results compared with the known ground truth. **(A)** To measure the effect of different signal-to-noise ratios, images were generated with three different levels of signal and noise intensities (100 each), showing that a critical SNR value of at least 4 is required to extract the most from the data in this assay. **(B)** To measure the effect of network density, images were generated with 5, 10 and 15 filaments (100 images for each), showing increased network density reduces sensitivity. **(C)** To explore the effect of the threshold parameter, four different threshold values were tested (100 images for each), highlighting the trade off between sensitivity and precision and that the optimal range is 87-92. **(D)** Finally, to investigate the effect of the numerical aperture (NA) of the lens, images were generated with a range of point-spread functions (100 for each), showing that even with high quality data, a minimum NA of 0.7-0.9 is required to achieve the resolution required.

Second, to test the effect of network density, a range of filament numbers (in a fixed size image) were considered. The noise value was chosen to be around the point where the algorithm performance changes rapidly, so that differences could be seen more easily. As shown in Fig. 4B, increased network density leads to a drop in sensitivity while precision remains relatively unchanged. This is because of the high default threshold value, chosen to ensure minimal noise is measured for high quality data. While a reduction in the total number of filaments reduces the chances of filament crossover and close-proximity, it does not remove these possibilities, as can be seen for the precision outliers with only five filaments. A lower network density leads to a reduced network size and therefore any errors will constitute a larger percentage error.

Third, to measure the impact of the algorithm threshold parameter (the most sensitive parameter), this parameter value was varied from its default value of the 90^th^ percentile of the intensity distribution of the image. The results, shown in Fig. 4C, highlight the trade off between sensitivity and precision. A higher threshold ensures greater precision, but at the cost of sensitivity. Conversely, dropping the threshold increases sensitivity, but beyond a point the precision falls and noise begins to adversely effect the output. Depending on the structures of interest, the resolution available and the experimental assay used, this value will need to be adjusted by the user to achieve the desired results.

Finally, to test the effect of the microscopy assay and resolution, the point spread function was varied to mimic various numerical apertures (NA) relevant to confocal microscopy. The results, shown in Fig. 4D, were tested using a high signal-to-noise ratio as all other tests were done on the point spread function mimicking the high NA lens setup, allowing the changes purely due to resolution to be clearly seen. For high quality data, we found a critical NA of 0.7-0.9, where the resolution becomes too poor and the sensitivity of the algorithm suddenly decreases. When designing and testing a microscopy assay, it is important to know where this limitation lies in order to ensure the extraction algorithm performs well. This may be particularly important for light-sheet microscopy where NA is limited by the physical configuration of the modality.

For high quality data, the sensitivity only drops when multiple filaments cross or are in close enough proximity that they begin to blur together and the tube-enhancing filter cannot distinguish them. The filaments here are drawn as 5nm thick and then blurred to match the resolution of our microscopy assay. Any filaments that fall within about 200nm of each other are likely to be nearly indistinguishable in accordance with the Rayleigh criterion. This also means that any crossings, even those that are completely perpendicular, are likely to be seen as globular instead of filamentous and may therefore drop below the threshold. In the cases where they do not, the lack of resolution and the skeletonisation process will represent the crossing poorly, typically appearing more as an ‘h’ or ’x-wing’ shape (see SI for details).

In order to assess algorithm suitability to intermediate filaments and microtubules, we also tested the effect of filament thickness. We used supersampling to create filaments of width 5nm (F-actin), 15nm (intermediate filaments) and 25nm (microtubules) and downscaled to fit a 95nm^2^ pixel size (which matches our microscope assay). No differences were found in the success metrics of our algorithm, as expected for objects significantly smaller than the available resolution. This demonstrates that our algorithm is likely to be easily adaptable to deal with other elements of the cytoskeleton.

### 2.4 Application 1: *Arabidopsis* Hypocotyl Cells

Wild-type (Col-0) *Arabidopsis* hypocotyl epidermal cells with GFP-Lifeact were imaged and the actin network extracted as described above. These were compared to cells with loss-of-function mutations either in Arp2[76] or in three formin genes (Formins 4, 7 and 8[44]). These were chosen as they are both mutants of actin filament nucleation regulators with stress-response phenotypes. First, a MANOVA (multivariate analysis of variance) test was used to determine if significant differences existed based on eleven dependent variables (skeleton density, branch point density, 2D filament angle, structure size and number density, filament width, deviation, signed and unsigned curvature, branch ratio and average 3D filament length). Only these eleven measures were used, rather than the full set of seventeen, because the unused measures were either unimportant (cell orientation), used in other measures (cell size, filament angle from imaging plane), redundant (2D equivalents of the 3D measurements) or unsuitable for this dataset (branching angles are not defined if no branches are detected). The MANOVA test uses a linear combination of the dependent variables to create a single combination variable that is then tested against the genotypes. This showed significant differences between the three genotypes (*p* < 0.001), although it cannot reveal exactly where these differences lie. Next, to determine the variables most responsible for this significant difference, the Tukey honest squares difference (HSD) post hoc test was used, which identified a number of key differences between the wild-type, *arp2-1* and *formin4/7/8* plants.

First, a statistically significant difference was found in the density of cytoskeletal structures (see Fig. 5A). While the *arp2-1* knockout showed no significant difference to either the wild-type or the formin mutant, the *formin4/7/8* mutant displayed a statistically-significantly increased structure density compared to the wild type (*p* = 0.046), with an average increase in density of 31%. Given that no differences were found in the average size of these structures, this suggests that the formin mutant cells may have more filamentous actin, leading to a higher network density.

**Figure 5:**
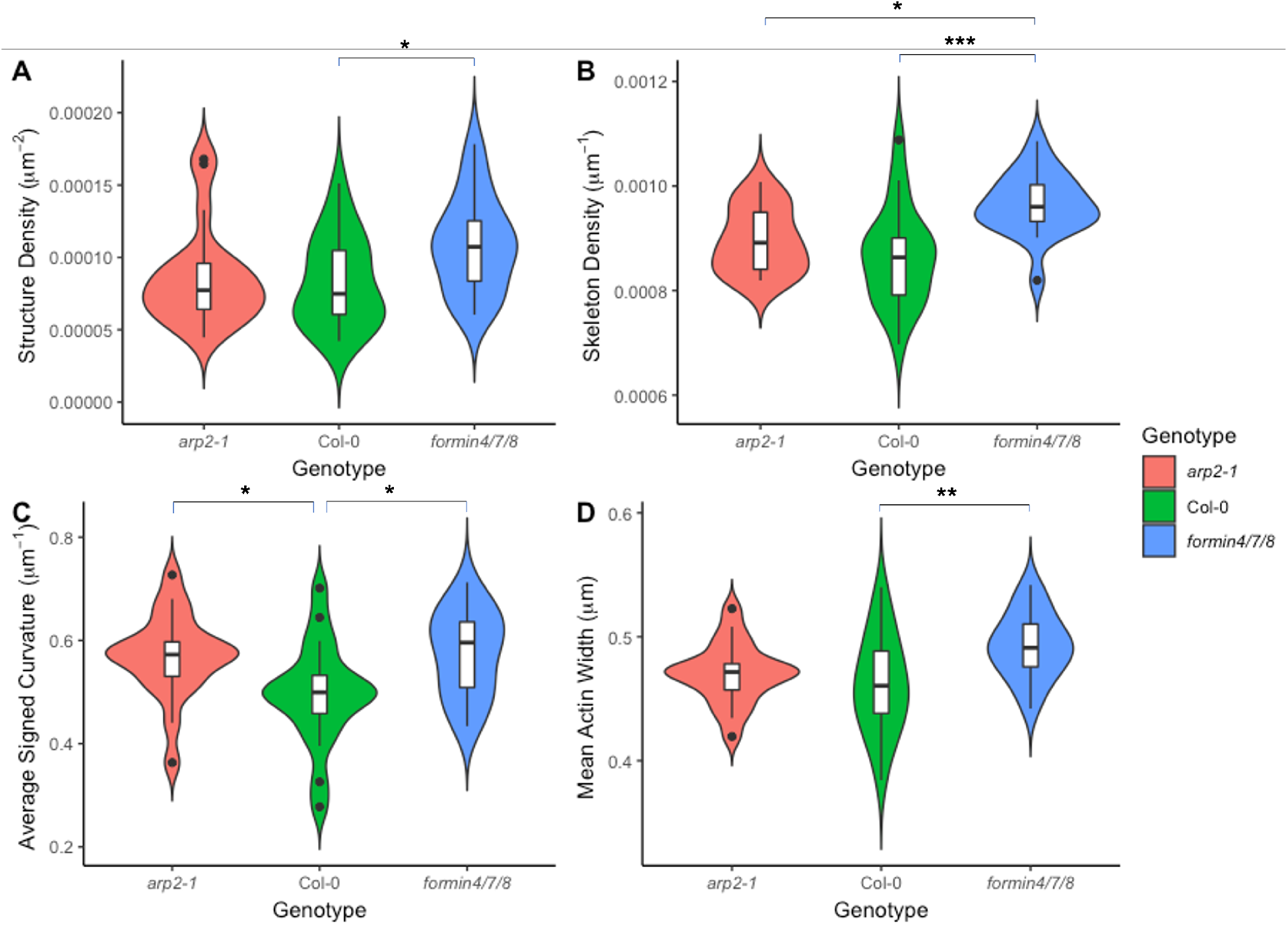
Comparison of the actin network in hypocotyls of the wild type, formin mutant and Arp2 mutant. Four measures (structure density, skeleton density, average signed curvature and mean actin width) showed significant changes between the wild type and *formin4/7/8* triple mutant. One of these (average signed curvature) also showed a significant difference between the wild type and *arp2-1* mutant and another (skeleton density) displayed a significant difference between the *arp2-1* and *formin4/7/8* mutants. Pairwise testing was performed with Tukey HSD after a MANOVA test. Single/double/triple asterisks represents *p* < 0.05, 0.01, 0.001 respectively.

Second, the skeleton density was also significantly greater in the *formin4/7/8* mutant compared to both the wild type (*p* < 0.001) and the *arp2-1* mutant (*p* = 0.012), while the *arp2-1* mutant was similar to the wild type (see Fig. 5B). This again suggests a more tightly packed network in *formin4/7/8* deficient cells, perhaps due either to increased filamentous actin content or increased cytoplasmic volume per unit cell surface area. A reduction in coupling of the network to the cell membrane is unlikely to significantly reduce the cell surface tension, however, due to its relatively small contribution compared to the internal turgor pressure.

Third, compared to the wild type, the average signed curvature of filaments was increased in both the *arp2-1 (p* = 0.029) and *formin4/7/8* (*p* = 0.020) knockouts (see Fig. 5C). As the Arp2/3 complex and formins are involved in coupling the cytoskeletal network to the membrane, a reduction in coupling could result in a drop in filament tension. It is plausible that this tension creates more linear filaments, so that a reduction in tension would lead to more curved bundles in the mutants.

Finally, the mean width of actin bundles was higher in the *formin4/7/8* knockout compared to the wild type (*p* = 0.0075) whereas no statistical significance was found with the *arp2-1* mutant (see Fig. 5D). While the difference in thickness is smaller than our pixel size (32nm versus 95nm), this is an average over filaments, which are then averaged over their length, meaning that small differences can be detected. This may not seem surprising given the greater network density measured in *formin4/7/8* knockout cells, but some formins have been shown to have a function related to bundling and thicker bundles tend to be present in hyper-elongated cells, whereas the *formin4/7/8* mutant phenotype in hypocotyls showed slower elongation and smaller cells. This suggests that these cells may already be adapting in order to try to reduce these deficiencies.

### 2.5 Application 2: Comparing Hypocotyl and Leaf Cells

We next examined the actin cytoskeleton in epidermal cells across two tissues: hypocotyl and leaves. To do this, all hypocotyl experiments described above were repeated on 4-6-week-old *Arabidopsis* leaves, their actin networks were extracted and then compared. The MANOVA test was used again, this time with an additional independent variable (cell type). Inclusion of leaf cells rendered the combined tissue genotype less statistically significant (*p* = 0.056), whereas the cell type itself showed significant differences (*p* < 0.001). Hypocotyls and leaves have epidermal cells with distinct morphologies and different rates of cell expansion, which may be the dominant reason for differing cytoskeletal characteristics. For example, hypocotyl cells exhibit accelerated directional growth (these cells must rapidly expand and elongate to reach the soil surface in order to have access to light). This rapid growth needs to be supported by the cytoskeleton[18] and therefore a lack of important actin binding proteins may impact this growth rate and result in more exaggerated differences between the genotypes than in the hypocotyl cells.

Given this MANOVA result, the Tukey HSD post-hoc tests were then performed on only the wild-type data in order to minimise any confounding variables. All genotypes are shown in Fig. 6 for completeness, which highlights the subtlety of the phenotype in the mutants compared to the significant differences between cell type.

**Figure 6:**
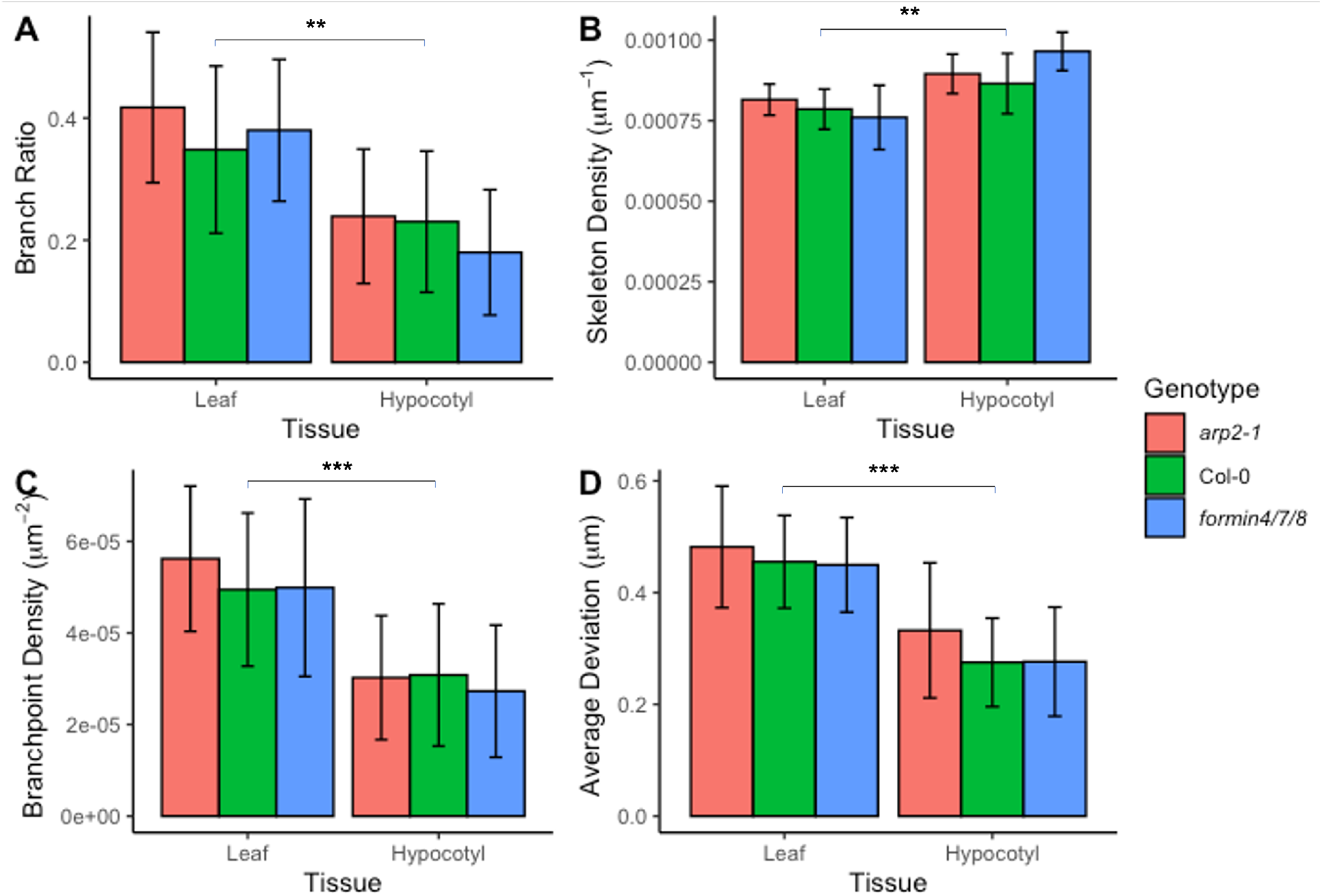
Effect of cell tissue on actin cytoskeleton. Four measures showed significant differences between wild-type hypocotyl and leaf cells. Significant differences are only shown between wild-type cells, although similar significance values were found between most genotype pairs. Pairwise testing performed with a Tukey HSD test after a MANOVA test. A double asterisk represents *p* < 0.01 and a triple asterisk denotes *p* < 0.001. Error bars show one standard deviation from the mean.

The branch ratio and branch point density, shown in Fig. 6A and Fig. 6C respectively, were both significantly higher in wild-type leaf cells than hypocotyls (*p* = 0.0043 and *p* < 0.001 respectively). The increased branching in leaves may be partly due to the less restrictive nature of the cell shape, allowing the network to form crossover junctions closer to perpendicular angles. This is likely to also be influenced by the large levels of cytoplasmic streaming that are observed in the long, thin hypocotyl cells. This streaming is generated through myosin motors on the actin filaments and bundles. Because the direction of motor assisted transport is defined by the structure of the cytoskeleton, these filaments are mostly aligned in the direction of the cytoplasmic flow[77].

The skeleton density of wild-type leaf cells was lower in leaf cells compared to hypocotyl cells (*p* = 0.0020), shown in Fig. 6B. While the *formin4/7/8* knockout was significantly different to the wild type in hypocotyls, the leaf cells appear much more consistent across genotypes. This difference in tissues is again likely down to the different cell shape and size constraints: perhaps more space afforded by wider cells and a smaller vacuole, as well as less reliance on cytoplasmic streaming, reduces the need for such an organised, polarised network in leaves.

Finally, the average deviation of filaments and bundles from linear is significantly different (*p* < 0.001) between cell types, with greater deviation in leaf cells compared to hypocotyls (see Fig. 6D). As with the other differences, this is possibly due to a different set of requirements for the cytoskeletal network due to differences in cell shape and developmental stage. For optimal cytoplasmic streaming in the long and narrow hypocotyls, the actin network needs to be highly parallel to the major cell axis and straight. This is in contrast to the highly irregular shapes of leaf cells, which have a width much closer to their length. The curved cell membrane and cell wall impart significant deviations in the linearity of actin filaments near the cell edges.

### 2.6 Application 3: *Blumeria* Infections

Wild type, *arp2-1* and *formin4/7/8* knockout leaves were all infected with *Blumeria graminis* (*Bgh*) and imaged at 48 hours post infection. MANOVA testing showed no significant changes due to genotype (*p* = 0.25), pathogen (*p* = 0.079), or a combination of the two (*p* = 0.086). Despite this, it is possible that some combination of measures could still show a significant difference. To test this, a principal component analysis (PCA) was performed, which scales, shifts and combines all variables into an alternative, yet equivalent, set of principal components. The aim of this is to encapsulate as much of the variation in the data as possible using only a subset of the principal components. The variance as a function of the principal component number is shown in Fig. 7, with the dotted line showing the mean percentage of variance over all principal components. The first four principal components contribute more than this mean, containing between them 77% of the total variation. By selecting only these four components and performing another MANOVA test, genotype (*p* = 0.037), pathogen (*p* = 0.0020) and their combination (*p* = 0.033) were all significant.

**Figure 7:**
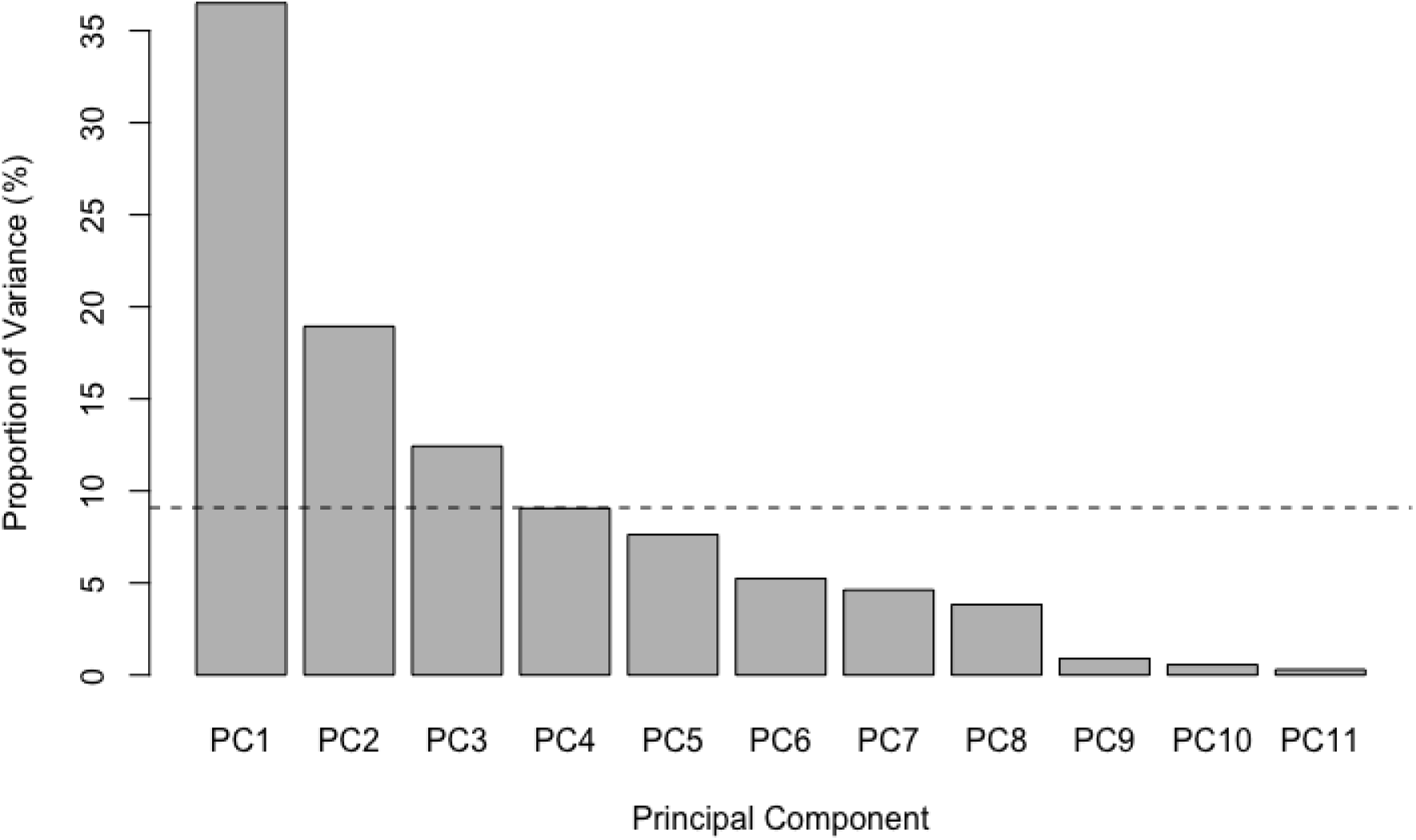
Principal component analysis (PCA) of *Blumeria graminis*-infected and uninfected leaves. Shown is the percentage of the variance that is explained by each of the principal components. The dotted line shows the mean percentage of variance over all principal components. The first four components contribute more than or equal to this mean and are therefore kept for future analysis.

*Bgh* was chosen as a pathogen due to its reported ability to perturb the actin network and direct trafficking during non-host immune responses[22]. Cell wall reinforcement and successful defence have been shown to be dependent upon actin dynamics[78]. These effects are however highly localised and their impact upon the overall properties of the cytoskeleton across the cell is likely to be subtle. However, it is promising that, despite this, our algorithm is still able to identify significant differences between infected and non-infected cells (albeit without allocating these to specific descriptors). With a larger sample size or higher resolution images, a better understanding of the filamentous actin around the infection site would likely be achievable by using our algorithm at the sub-cellular level to compare local network properties.

## 3 Discussion

In this study we have developed a novel software algorithm, the DRAGoN tool, that can automatically extract and quantify multiple characteristics of complex actin networks, with particular focus on plant cells. Our software is designed to work with both 2D and 3D data. We have demonstrated its utility by establishing that it can accurately extract the large-scale actin structure for a range of cell types and mutants.

Our tool provides seventeen distinct quantitative measures of actin networks (see Table 1), including properties of the overall network (such as skeleton density and branch ratio), individual filament properties (such as filament widths, lengths, directions and curvatures) and branching properties (such as branch angles). The code can easily be extended to include additional measures as required by the user.

We validated our algorithm in a number of ways. This involved generating artificial networks (where the ground truth was perfectly known) consisting of individual filaments with various properties, including a number of edge cases. The length, curvature and deviation of these filaments were measured with the algorithm and shown to have high (>90%) precision and sensitivity with parameters corresponding to typical imaging modalities. This process was then repeated with various branch angles, again with similarly high detection rates.

We then applied our code to several real examples from *Arabidopsis thaliana*. First, we examined how the actin cytoskeleton differed in wild-type cells compared to two null mutants: *arp2-1* and *formin4/7/8*. We found that, in hypocotyl cells, both mutants showed significant differences relative to the wild type in four separate measures, including filament curvature and width. The signed curvature was greater in both mutants compared to the wild type, while the average actin width was larger in the *formin4/7/8* mutant than the wild type. Both mutants had a higher skeleton density than the wild type, but only *formin4/7/8* had an increased structure density. This possibly points to the reduction of membrane anchoring points for the cytoskeleton, one function of formins[79], reducing tension and support in the network, which may result in adaptions being made in order to bolster strength.

Next, we examined hypocotyl versus leaf cells, finding significant difference in branching (leaves have a higher branch point density) and network density (hypocotyls have higher density). The detection of these significant differences between tissues highlights the different requirements of leaf and hypocotyl cells. It is plausible that the difference arises since hypocotyl cells require much of the network to run parallel to the cell’s long axis to aid with transport, increasing packing density of the cables and reducing the need for branching. This may further be reinforced by reduced deviation of the filaments in hypocotyl cells.

Finally, we examined the response of *Arabidopsis* leaf cells to infection by *Blumeria*. Identifying significant differences were more difficult in this case, although was possible by using PCA analysis. By taking the four principle components that contributed more than the mean total variance, we showed that *Bgh* has a detectable impact on the actin network, although we were unable to identify the particular microscopic network differences at play. Our analysis only considered the actin network of the whole cell. Since *Bgh* infection may only modify the cytoskeletal network close to the point of infection, we may have missed local network changes. This could be remedied by optimising data acquisition to analyse the network of the region around the infection site. This would need to be performed at diffraction-limited resolution and at high time resolution to prevent dynamic filament smearing in projected images.

The ability to identify differences in specific parameters can indicate the action of specific classes of actin binding proteins and their associated signalling pathways in tissue differentiation. The measured changes across genotypes shows that even more subtle differences can be detected and measured, which should allow even better insight into the mechanisms underlying the cytoskeletal processes. While we were unable to quantify the differences in the network induced by pathogen invasion, the fact a difference could be found using principal component analysis suggests that as yet undefined ratios of network properties characterise the response to pathogen assault. The changes in proximity to the appressorium are often highly visible, whereas the rest of the network appears to remain similar. It is possible that changes occur here, but they will be very subtle, therefore most of the changes are going to occur in a small region and measurements across the whole cell will only change by a very small amount. A much larger data set or perhaps an artificial stimulation of the immune response (e.g. a microneedle assay[80]) may help in discerning these changes in more detail.

To facilitate further development or optimisation for particular data sets, we have made the DRAGoN software freely available and open source at https://github.com/JordanHembrow5/DRAGoN. The flexibility and non-specificity of this tool is one of its main advantages and should enable it to be useful in a range of organisms, mutants, tissues, cell types and environments. A number of key parameters (particularly those for the filtering and skeletonisation steps) can be adjusted to best fit a given image modality and labelling method.

The noisy nature of both the actin cytoskeleton and microscopy means that a perfect network extraction tool cannot exist. Our tool has a number of limitations. First, we focus on extracting actin bundles rather than the finer F-actin structure (which was motivated by the nature of our training data but precluded analysis of highly dynamic filament populations). Second, while the code has support for 3D data sets and measurements, these are currently implemented in only a simple manner by estimating z-positions from stack intensities. In future, better isolation of the filament position in 3D would enable additional information to be gathered on measurements such as curvature and branch angles, as well as enable new types of 3D measurements. Third, our tool is designed to work at single time points. A future extension could involve frame-by-frame filament tracking. This would allow network remodelling to be analysed in greater detail, and the network in nearby time points could be used to inform and improve filament detection, providing more information than is possible by using single images in isolation.

While the software was designed using data from *A. thaliana*, particularly images from leaf and hypocotyl cells, we have aimed to create a general resource that is easily adaptable to other cell types and organisms. For example, quantifying the network structure in other plant tissues and cell types, such as roots and root tips, would likely be straightforward. Further, we expect our algorithm (with suitable parameter changes) will also perform well on other organisms, including animal cells. Structures other than actin are likely to be analysable with only minor modifications. For example, extracting the microtubule network is a substantially easy task than that for actin due to their increased and more-consistent thickness. However, it is worth pointing out that there will be significantly more crossing events for microtubules and so an improved method of network reconstruction (perhaps by more fully utilising every image in a 3D stack) would be needed. Finally, it will be worth investigating whether out approach can be used to extract multiple different networks from single images. For example, since filament width is already on output of our algorithm, it may be relatively straightforward to add a method that segments the cytoskeleton into F-actin, intermediate filaments and microtubules (given sufficient labelling and resolution to distinguish between them).

In addition to the points mentioned above, there are a number of further ways that our tool could be extended in future. First, we have used the 3D data provided by z-stacks only to estimate 3D filament lengths and angles. More fully reconstructing the full 3D network could have a number of important applications, particularly in animal cells where the lack of a central vacuole increases the importance of the full three-dimensional actin structure. Second, we currently analyse images at single time points. Identifying the network at neighbouring time points (with, for example, some modified Hungarian algorithm) could lead to both an improved accuracy of network extraction and a way to quantify dynamic network changes (for example, in response to fungal attack). Third, previous work in this area utilised analysis of the actin structure to measure organelle motility[47]. With our algorithm, this could naturally be extended by using the improved network quantification that our software yields. By including measurements such as curvature, matching organelle movement to actin tracks is likely to be more accurate and provide additional data to explore mechanisms and design mathematical models. Fourth, since analysis of a single frame takes only a couple of seconds with our algorithm, network extraction could be performed in real time, with results instantly fed back to the microscope user. Near real-time analysis could inform which areas of the cell to probe for an immune response or for how long to apply a perturbation (such as in the nanoindentation experiments that have been used to test for actin remodelling to a physical stimulus[23]). Further, for laser dissection experiments, immediate and detailed information and statistics of the actin network could suggest which cables to sever.

Our DRAGoN tool demonstrates that accurate extraction of the actin network structure can be accomplished with only minimal human interaction. Further, the same general algorithm will be relevant across cell types, tissues, organisms and network types. Applications of this work are widespread and range from basic biological understanding to global crop security. As a result it is likely that such tools will become increasing common and important in biology as improved methods of imaging the fine detail of actin and other cytoskeletal elements become available.

## 4 Methods

### 4.1 Image Analysis Algorithm

We break our network extraction algorithm into a number of sequential stages—initial steps, skeletonisation, image rotation, skeleton labelling and measuring properties—which we discuss in turn. For full details of our method, see the Supporting Information.

#### 4.1.1 Initial steps

The initial step involves background filtering. While a Gaussian blur or median filter can smooth and modulate noise, they do not preserve edges[81], which are critical in filament detection. Conversely certain non-linear techniques, such as the rolling ball algorithm, are able to preserve edges and features whilst removing noise and non-linear illumination[82]. Therefore, based on a rolling ball method, our raw images are first filtered to remove background noise and uneven illumination through a top-hat filter with a sphere shaped structuring element (via the *imtophat* function in MATLAB). The disk should be wider than the structures that must be preserved. However, since the algorithm time roughly scales with the square of the disk radius, there is a necessary balance between accuracy and speed. A radius of 15 pixels (approximately 1.4μm in our images) was chosen as optimal for the actin structures we encountered.

#### 4.1.2 Skeletonisation

The resultant image then has fibrous structures enhanced by using a Hessian-based multi-scale filter (via the *fibermetric* function in MATLAB). No thickness values were passed to this filter as actin bundles vary in thickness and we wish to preserve all of them. The resulting significant contrast between filaments and the background then allowed a simple threshold to produce a binary image. We found a value of the 90^th^ percentile of the fluorescence intensity produced reliable results for our data. Any elements containing fewer than 20 pixels is then removed, and the binary was eroded by the *bwskel* function, which removes pixels from the perimeter of objects until a single pixel thick skeleton remains.

#### 4.1.3 Image rotation

Images are then rotated so that the long axis of the cell aligned with the horizontal axis. We tried rotating both before and after the skeletonisation step, finding that rotation first can lose some information due to the discrete nature of the pixels, while rotating the skeleton afterwards can result in fragmentation of areas. In the end, we found the best result involved performing the image rotation both before and after the skeletonisation process, with the results then combined with pixel-wise OR logic. The results are then eroded away again to ensure each filament is only one pixel thick.

#### 4.1.4 Skeleton Labelling

We label each filament in the skeleton with its own unique integer, such that only the pixels within that filament have that number. To avoid connected branches having the same label, we first temporarily remove the branch points using the *bwmorph* MATLAB function. Next, we flood-fill all the components of the network to label the different isolated filaments, using a combination of *bwconncomp* and *labelmatrix*. A side effect of this process is that long filaments with multiple branches are broken into distinct pieces between branch points. To rectify this we use a custom joining algorithm to re-join filaments (see Supporting Information for details) so that, at each branch point, the two filaments that are closest in direction are joined. This is done for every branch point and a final pass ensured that the labelling is continuous for all filaments.

#### 4.1.5 Measuring Properties

After every filament is uniquely labelled along with a list of start and end points, a number of quantitative measures can be calculated. First, filament length in two dimensions is found by counting the number of pixels with a given filament label with diagonal connections weighted by an additional factor 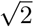 (see SI for details). If three-dimensional data (*i.e*. multiple z-stacks) is available, the filament length is extended to three dimensions. To do this, the location of the brightest pixel in the image stack is used to determine the z-position at each point in the skeleton, from which the mean angular displacement of the filament relative to the image plane can be calculated. The 2D projection length then gives an estimate of the real 3D length (see SI).

Second, filament width is estimated by measuring the local direction of a filament and then measuring the length of the binary image (before erosion for skeletonisation) perpendicular to that direction. This is repeated for every pixel along the length of the filament and averaged to find the mean filament width.

Third, filament angles are defined relative to the major axis of the cell. Using the MATLAB *regionprops* function, the binary mask that was initially used to remove all data outside of the cell can be used to determine the cell orientation relative to the horizontal axis. The image can then be rotated such that the major axis is parallel to the horizontal axis. This is not necessary for cells without any polarity. Once the image is rotated, all labelled filaments have their 2D orientation determined by finding the displacement in *x* and *y* between their endpoints and using the inverse tangent. As no information is typically available about the direction of the individual filaments, their angle is only defined in the range 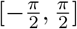. The same can be repeated in 3D by calculating the displacement and therefore the angle relative to the image plane.

Fourth, filament curvature is defined using the Menger curvature: the inverse of the radius of the circle that passes through three points[83]. Each internal pixel (no curvature is defined at the endpoints) in a filament is chosen as a central point. Then the adjacent points a characteristic length λ away are determined. For pixels closer than λ to an endpoint, the largest possible distance is chosen (*i.e*. the endpoint itself is used). The curvature is measured for these three pixels, iterating along the filament and averaging to determine the average curvature of a filament. The deviation of a filament is defined as the average distance between each pixel in the filament and the straight line joining the two endpoints.

Finally, calculation of the branch angle requires a branch point and two or three labelled filaments connected to it. The filament end points not connected to the branch point are determined and used in conjunction with the branch point itself. Each unique pair of end points (three in total) are combined with the branch point in the middle to produce three different angle pairs. The angles with the smallest deviation travelling from one filament to the branch was taken to be the main filament, with the middle angle then defined as the branch angle. The branch ratio is a measure of how much a network system branches, defined as the total number of branch points divided by the number of individual filaments.

All of the available measures can be found in table 1

### 4.2 Plant and Pathogen Growth

#### 4.2.1 Plant Material

*Arabidopsis thaliana* ecotype Colombia-0 (Col-0) with GFP-Lifeact were grown as the wild type, alongside loss-of-function mutations in Arp2 (SALK SALK_003448) or three formin genes (Formins 4, 7 and 8) as described in Sassmann et al[44].

#### 4.2.2 *Arabidopsis* Hypocotyls

*A. thaliana* seeds were sterilised with Cl2 for 4-5 hours in a sealed container by mixing 100ml of bleach with 3ml of 37% HCl. Sterilised seeds were suspended in molecular biology grade water and stored in the dark at 3°C for a minimum of 5 days. To produce the extended hypocotyls, seeds were dark grown in a humid growth chamber at 21°C for 5 days. 100μl half-concentration Murashige and Skoog (MS) growth medium with Gamborg’s Vitamins containing 0.8% w/v agar was used as the growth medium, upon which the seeds were placed. 500μl centrifuge tubes were used to contain the medium and support the hypocotyls.

#### 4.2.3 *Arabidopsis* Leaves

10-15 *A. thaliana* seeds were sown onto F2 soil with sand, mixed with vermiculite in a 3:1 ratio. The seeds were left in the dark at 3-5°C for a week, then transferred to a growth cabinet (16h light, 8h dark) at 21°C. After 1-2 weeks more, plants were transferred to a new pot as to grow uncontested.

#### 4.2.4 Blumeria Graminis

*Blumeria graminis* (*Bgh*) was cultivated on “Golden Promise” barley (16h light, 8h dark) at 17°C by weekly infection of three-week old barley plants.

#### 4.2.5 *Blumeria* Infection

After 7 days of cultivation on barley, *Bgh* spores were sprinkled on whole plants for the elongated hypocotyls; they were placed back in the dark at 17°C for 24 hours and remained in their centrifuge tubes. For leaf infections, 4-6 week old A. *thaliana* plants were cut at the base of the petiole and placed on damp filter paper in a petri dish before spores were sprinkled on the adaxial (dorsal) surface. This assay was left in the dark for 48 hours at 17°C before imaging.

### 4.3 Imaging Methods

Images were taken with a spinning disc confocal microscope, equipped with a 60x lens and NA of 1.35. A laser power of 7mW at a wavelength 488nm was used for imaging GFP-Lifeact. Images were taken with 200ms exposure and z-stacks had a separation of 0.55μm. Entire hypocotyls were mounted while 5×5mm squares of leaf were cut, away from the central vascular tissue, in order to keep everything as flat as possible.

## Acknowledgements

JH, MJD and DMR acknowledge that this work was supported by the Biotechnology and Biological Sciences Research Council-funded South West Biosciences Doctoral Training Partnership [BB/M009122/1]. MJD and DMR were also supported by a Wellcome Trust Institutional Strategic Support Award (WT105618MA). DMR gratefully acknowledges financial support from the Medical Research Council (MR/P022405/1).

## Notes

https://github.com/JordanHembrow5/DRAGoN

### Competing Interest Statement

The authors have declared no competing interest.

