## Supplementary Text for "AUTOMATIC EXTRACTION OF ACTIN NETWORKS IN PLANTS"

### Automatic Extraction of Actin Networks in Plants: Supplementary Information

January 18, 2023

#### S1 Algorithm Design

Properties of individual filaments are determined by iterating through filament labels and creating a binary image of each filament. Label numbers are contiguous from 1 to the number of filaments, so the filaments are first separated into their own binary images by only keeping the pixels with the number equal to that of the filament label. This enables measurements to be made more easily, without interference with other filaments and allows the easy storage of data in an array, where the index corresponds to the filament label. This process allows the leverage of powerful tools built-in to Matlab, such as *bwmorph* for manipulating binary images and skeletons through a range of morphological operations.

##### S1.1 Filament Orientation

Filament orientations in the two-dimensional imaging plane, frequently referred to in the program as the XY plane, are relative to the major axis of the cell. The major axis is found by fitting an ellipse to the cell mask which has an angle from the x-axis to the major axis of the ellipse with the same second-moments as the cell mask, via the *regionprops* function. The image is then rotated by this angle such that the major axis is now aligned with the x-axis, allowing easier comparisons between cells. Filament orientation doesn't use *regionprops* however, but instead takes the line joining the end points, which are found via *bwmorph*, and calculates the angle relative to the x-axis via the inverse of the tangent function. This ensures the ultimate direction of cargo travelling along the network is measured, despite any kinks in the filament due to microtubule movement or long curves around organelles or even curved walls and membranes. As the 2D image used for extracting the network is a maximum projection of a stack of 2D images at a range of $z$-positions, this stack is used to estimate the 3D position of the filaments. For each pixel in the skeleton, the intensity of the corresponding pixel in each image of the stack is measured and compared, with the plane of the greatest intensity being reported. While resolution in $z$ is much poorer than in the imaging plane, an estimate of the filament orientation can be made based upon stack separation and these maximum pixel values. The cosine rule is used to estimate the angle of the filament in 3D relative to the filament in the maximum projection.

##### S1.2 Filament Labelling

Correctly labelling each filament enabled all of the measurements made in the analysis tool. This process was done in stages, further refining the labelling at each stage to accurately represent the network. Fragmentation of the network was the first step, to give each filament, between branch points, and on branches, a unique label via a flood-fill algorithm. This was achieved by determining the branch points with *bwmorph* and removing them from the skeleton. Multiple passes had to be made to ensure that filaments connected horizontally or vertically weren't still connected diagonally after the initial branch point removal. As this process split filaments at every branch point, these points were checked in order to determine how to rejoin the network with the branch getting one label and the main filament another. Comparing the orientations of different filament pairs connected to the branch point allowed the straightest pairing to be giving a single label and rejoined, while the branch was left with a different label. A parameter for the characteristic length

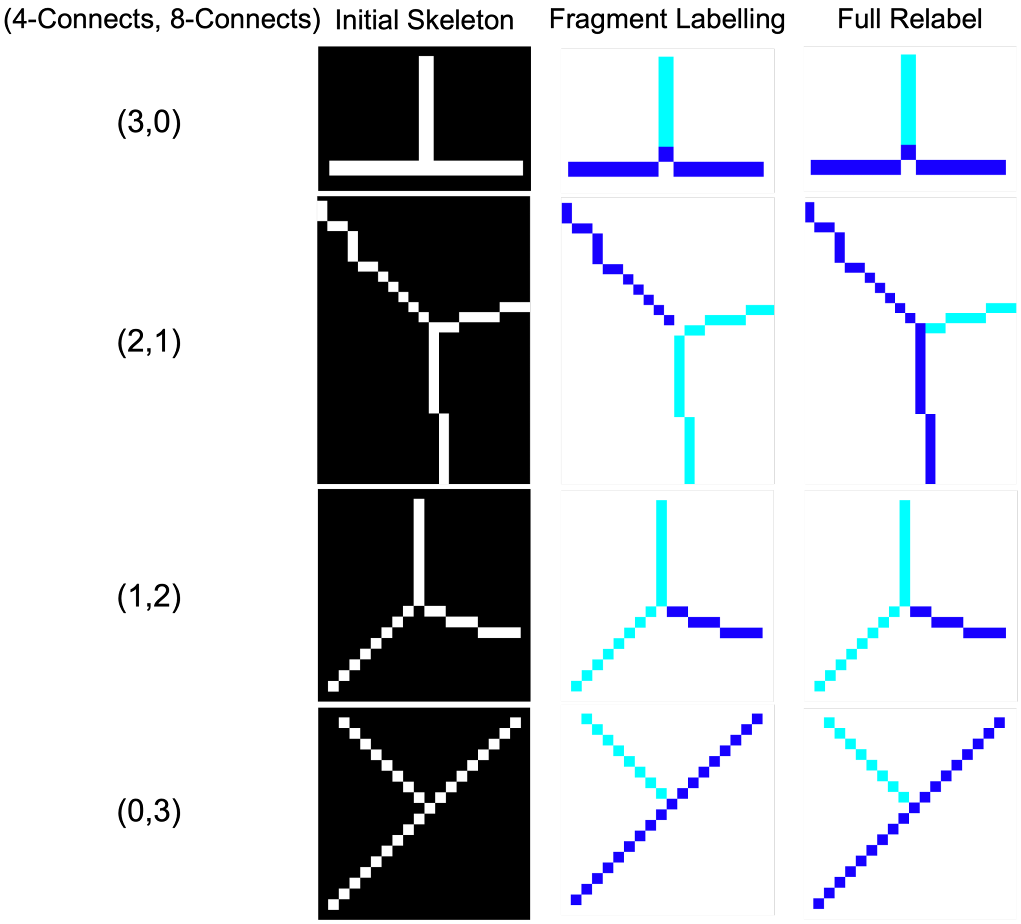


Figure S1: **Filament Labelling:** Flood fill algorithms used for labelling filaments require disconnects at branches to function as desired, otherwise every pixel in the skeletons shown would have the same label, making the process redundant. This fragmentation process can cause issues, however, as remaining 8-connected filaments can connect the network incorrectly in some cases. Additional checks and relabelling have been implemented in the right-most column to address this infrequent issue.

away from a branch point was used for the filament orientation, as long cables are free to bend and change orientation far from the branch and this shouldn't influence the choices made. As this process resulted in a label number being unused, these numbers were stored and once the process was complete, the highest number labels were replaced with these missing values. This ensures that for $n$ filaments, the labels are always from 1 to $n$. In some cases, the branch point removal only resulted in two filament labels, but had the labelling still incorrect, as shown in figure S1. Relabelling these filaments is more problematic, and this only occurs when a branch point has two 4-connected pixels and one 8-connected pixel, albeit only some of the time. All of these cases are checked, with the 4-connected pixels removed in order to yield 3 fragments, with another flood fill process occurring to separate the labels. The same algorithm as described above can be used to determine the correct labelling, with the extra step of replacing those previously removed 4-connected pixels, and converting the new label numbers (always 1,2,3) back to whatever the original pair of labels were. This process only had two labels to begin with, so no renumbering was required.

#### S1.3 Filament Length

Filament lengths in the XY plane are determined by their number of pixels $N_{\mathrm{px}}$ and the way these pixels are connected. A fixed number of pixels orientated horizontally or diagonally will have a different length, and filaments which are not linear cannot be measured by calculating the distance between endpoints. The distance between 4-connected pixels (horizontally or vertically) is equal to the size of a pixel, but the distance between any diagonally connected pixels is larger by a factor of $\sqrt{2}$. To determine the number of diagonal connections, each filament is relabelled, but this time with a restriction that filaments can only be 4-connected. Therefore the number of labels $N_{\mathrm{lbl}}$ is equal to the number of diagonal connections plus one. From here, the filament length $l$ is calculated via equation S1.

$$l = N_{\mathrm{px}} + (N_{\mathrm{lbl}} - 1)(\sqrt{2} - 1) \tag{S1}$$

For 3D length calculations, the average angle of the filament relative to the imaging plane is first calculated (the 3D filament orientation), then the 2D length is scaled by the reciprocal of the cosine of that angle.

#### S1.4 Branch Angles

At a branch point, there are three possible pairs of filament directions that could be used to define branch angles: the angle of the main filament relative to itself before and after the branch as well as the angle between the branch and the main filament, both before and after the branch. If the main filament isn't perfectly straight, this gives three different angles. First, to measure the angles, a length scale is defined in order to avoid any kinks or deviations far from the branch in longer filaments. From here, the filaments are temporarily cut, and the three end points determined via *bwmorph*. The three ends form 3 unique pairs of ends, with the branch point being the third point of the triangle. These three angles are calculated via the cosine rule, and the middle angle is selected as the branch. This is because a small angle would be associated with going backwards down the main filament and having to change direction significantly to follow the branch, while a large angle is likely to be continuing down the main filament. The selected angle is then taken away from $\pi$ such that the final angle is actually a measure of the deviation a molecular motor would have to take in order to follow the branch instead of continuing on the main filament.

#### S1.5 Filament Width

Filament widths are determined by the average width at every point along a filament. For every pixel in a filament, the direction of the filament is found, such that the perpendicular direction can be calculated in order to determine the width. A parameter is used to determine the length scale over which to determine the direction, as longer lengths can be more accurate in the case of a straight filament but can be erroneous for curved filament, therefore a length of approximately 0.5µm was chosen to be suitable here. The binary image before skeletonisation is used to measure the width, as the filtering and thresholding process is consistent and provides a good estimate for filament width. The algorithm moves pixel-by-pixel along the width until the binary image yields a false value, then does the same in the negative direction, in case the skeleton isn't perfectly central. These two distances are added to give the width at this point, and this is repeated for every pixel to yield an average width.

##### S1.6 Curvature and Deviation

Almost all filaments in the networks analysed were not completely straight, so these deviations and the curvature needed to be characterised and measured. The deviation is a very simple measure in which a straight line was drawn between the filament endpoints, and the distance between this line and each pixel in the filament was measured, such that an average could be taken. The distance for the point $(p_x, p_y)$ away from the line defined by $(e_{1,x}, e_{1,y})$, $(e_{2,x}, e_{2,y})$ is given by

$$r = \frac{|e_{2,x} - e_{1,x})(e_{1,y} - p_y) - (e_{1,x} - p_x)(e_{2,y} - e_{1,y})|}{\sqrt{(e_{2,x} - e_{1,x})^2 + (e_{2,y} - e_{1,y})^2}}. \tag{S2}$$

The curvature of a filament, while always being defined as positive as it's the reciprocal of the radius, was made up of measurements of the curvature at every pixel, and therefore these individual components could be signed. Two separate measures are used here, where the components are summed as is, or the magnitude is always taken, before finding the mean and taking the absolute value of the final result. To measure the curvature at a given point, the position of the filament at a characteristic length $\lambda$ away from the point is chosen, in both directions. The triangle formed from these three points $(a, b, c)$ was used to measure the Menger curvature with

$$c(a, b, c) = \frac{1}{R} = \frac{4A}{|ab||bc||ca|}, \tag{S3}$$

where the triangle area is determined by

$$A(a, b, c) = \frac{1}{2}|(x_a - x_c)(y_b - y_a) - (x_a - x_b)(y_c - y_a)|, \tag{S4}$$

and the side lengths are

$$|ab| = \sqrt{(x_b - x_a)^2 + (y_b - y_a)^2}; \;\; |bc| = \sqrt{(x_c - x_b)^2 + (y_c - y_b)^2}; \;\; |ca| = \sqrt{(x_a - x_c)^2 + (y_a - y_c)^2}. \tag{S5}$$

The choice of $\lambda$ affects the result of the curvature due to the coastline paradox, and was used to uncouple the relationship between pixel size and curvature. The length defined here, is the maximum distance (always chosen if possible) between each point in the triangle. This process is repeated for all filament pixels not located at endpoints.

#### S2 Algorithm Validation

##### S2.1 Length, Curvature and Deviation

Figure S2 shows three filaments which contain the same number of pixels with substantially different lengths and curvatures. Each filament has its length measured, its average Menger curvature calculated, and the average distance from each pixel to the straight line joining endpoints determined. The curvature is determined on a length scale suitable to the resolution of the image and the capture device (up to 5px between sampled points in this case) and is calculated both in signed and unsigned variants, before the absolute value being taken. Filament a has the shortest length of 23 pixels, with filament b being the longest at 32.1 pixels and filament c is 27.1 pixels long. Filament a has no curvature or deviation, whereas filament b has the greatest unsigned curvature with a smaller deviation than c. For signed curvature, c has greater curvature due to its consistent direction relative to b.

Additional testing of curvature was performed on unit circles and alongside manual calculations for three given points, all of which were accurate.

##### S2.2 Branch Angles

A range of different angle cases were tested and are shown in figure S3. The scenarios were made to test a range of different edge cases, with straight lines between known coordinates of ends and branch points used to calculate the expected result accurately. As the algorithm uses a length scale over which to determine the angle, and the pixel network is discretised and therefore the lines are not perfectly represented, small errors

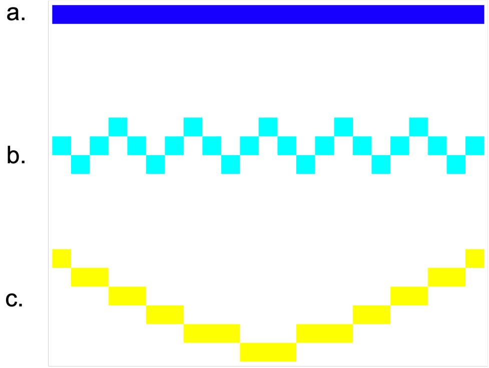


Figure S2: **Describing Filaments**: Various arrangements of 23 pixels represent filaments with different lengths, curvatures, and deviations from being linear. A range of statistics describing the different properties of these filaments is used to classify and differentiate them.

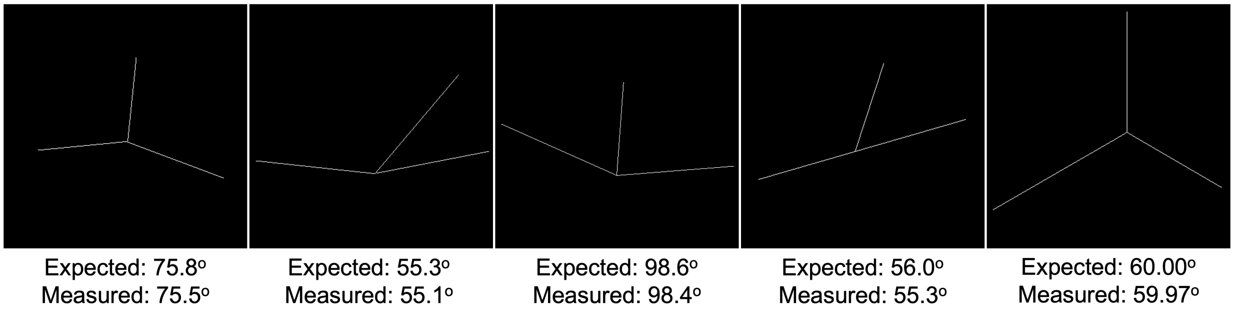


Figure S3: **Branch Angle Validation**: A range of different branch scenarios were tested, with known angles being compared to those measured. Small deviations from expected measurements are due to branch point estimates being a pixel out in some cases, or imperfect representation of lines onto discrete pixels, as well as small errors from not measuring the full filament length. Largest error is 1.25% and the smallest is 0.05%.

in the calculated angle are expected. The errors measured here range from 0.05% to 1.25% and therefore are perfectly valid in the scope of the project. The benefit of testing branch angles is that it encompasses testing filament angle measurements as well, and these must therefore also be accurate.

### S3 Skeletonisation Testing

#### S3.1 Superscaling

Sampling data more frequently (spatial, temporal or both) than the desired output and averaging over several data points to achieve higher resolution has been used for a wide range of applications. As the targeted microscopy assay for the testing has a pixel size of 95nm$^2$ and the filaments to be sampled started at 5nm thick, we employed this technique spatially when artificially generating filament data. To achieve the 5nm size, the pixels were cut into 19 subpixels in each dimension, resulting in a 19$^2$ increase in array size (200x200 -> 3800x3800). Any filaments were reconstructed in the new array with their positional co-ordinates being scaled by 19 and the desired thickness based upon filament width. From here a Gaussian blur with standard deviation matching the point-spread function of our microscope was applied to the array, with the pixel size scaled by 19 to match the new pixel size. From here, each 19x19 block of subpixels in the image was combined and averaged to reform the 200x200 image.

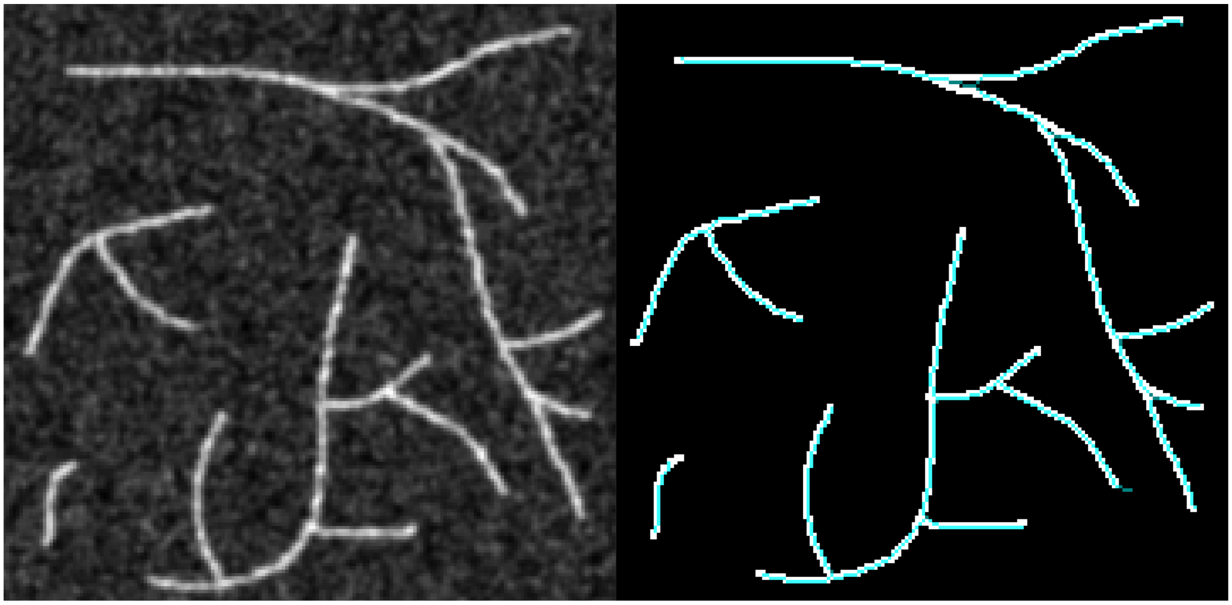

Figure S4: **Skeletonisation In An Artificial Data Set:** The initial image is shown on the left, with the skeleton that was extracted, roughly following the centre-line of the filaments, superimposed over the image on the right.

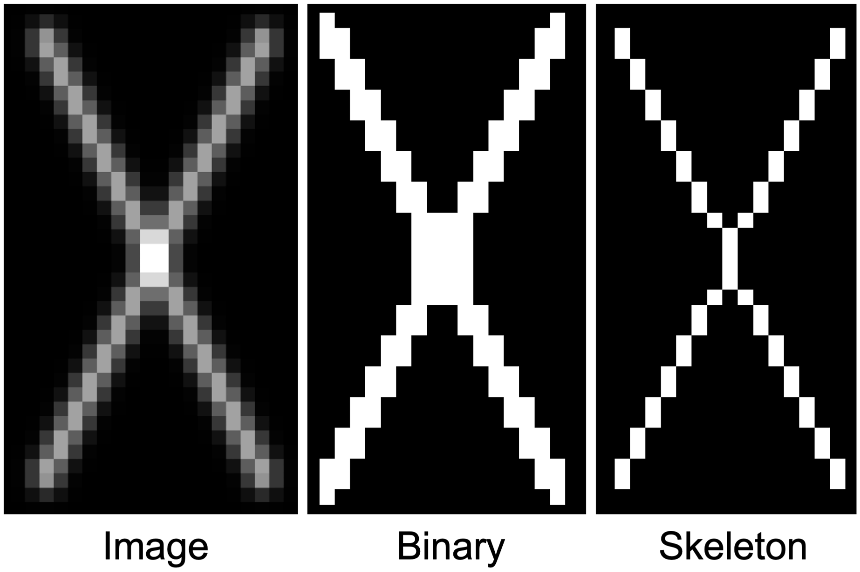


Figure S5: **Crossing Issues:** Two filaments intersecting frequently results in their skeletons combining for several pixels around their crossing. This makes evaluating branching and crossing very challenging.

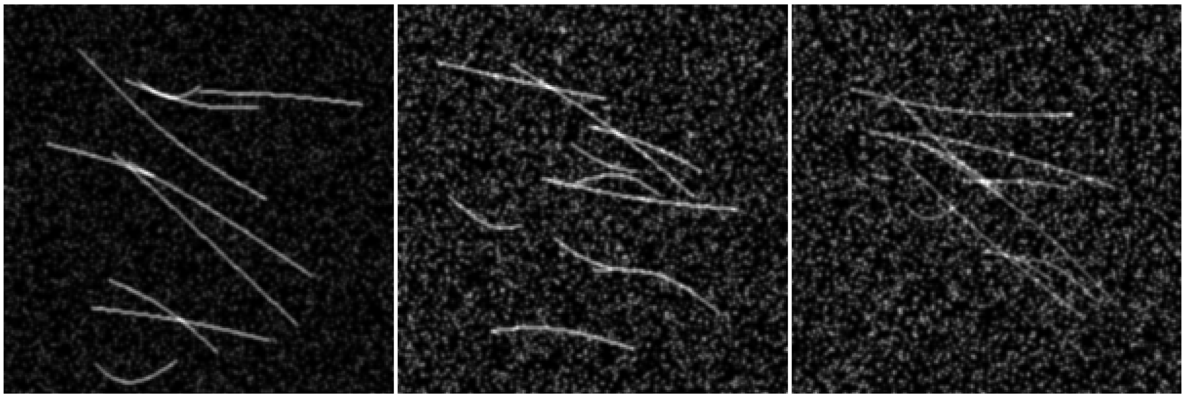


Figure S6: **Various SNR Levels:** Example images of artificially generated data at different SNR levels. As the SNR drops below 4, the amplitude of the noise eclipses the filament signal and any filaments in proximity of each other are indiscernible.

#### S3.2 SNR Variation

The images generated for skeletonisation testing were 200x200 unsigned 8-bit integer arrays (uint8, range: 0-255). Two random points in this array were chosen as the filament ends, by uniform random number generation of values between 1 and 200, for both co-ordinates. The midpoint of the line connecting these end points was calculated by taking their mean, and both the co-ordinates of this midpoint were shifted by a random amount between 0 and 15 to add curvature. 10 filaments were generated independently of each other and added to the image. These filaments were 3 pixels thick and had varying brightness depending on the signal-to-noise ratio being targeted. For high SNR, the filaments had an intensity of 80, while the medium and low SNR groups had an intensity of 60 and 50 respectively. A binary image of the filament-only image was also generated to give the mask for SNR calculations. To generate the noise, another 200x200 uint8 array was generated with salt and pepper noise, resulting in 40% of the values being 1 and the remaining 60% being 0. Each element of the array was scaled by a random number, ranging from 0 to the maximum noise value parameter, which varied depending on the SNR level being targeted. This was 80 at high SNR, with 120 and 140 being used for the middle and low SNR images. The noise array was added to the array containing the filaments, and a Gaussian blur (0.9 pixel radius) was applied to obtain the final image. The SNR is defined in equation S6. The mean value of the signal $\mu_{\text{sig}}$ was measured as the mean value of all the true pixels in the filament-only binary image, while the standard deviation of the noise $\sigma_{\text{noise}}$ was simply the standard deviation of all of the remaining pixels.

$$\text{SNR} = \frac{\mu_{\text{sig}}}{\sigma_{\text{noise}}} \tag{S6}$$

#### S3.3 Numerical Aperture

For a zero mean Gaussian distribution, we derive the full width at half maximum (FWHM) to determine the relationship between the minimum resolvable distance and the Gaussian standard deviation $\sigma$. A zero-mean Gaussian is given by

$$f(x) = \frac{1}{\sigma\sqrt{2\pi}} e^{-\frac{x^2}{2\sigma^2}}. \tag{S7}$$

If $x_{0.5}$ is the point that the Gaussian amplitude reduces to half it's maximum value, then the FWHM is defined as $2x_{0.5}$ due to the symmetry about the mean of zero, therefore we can write

$$f(x_{0.5}) = \frac{1}{2\sigma\sqrt{2\pi}} = \frac{1}{\sigma\sqrt{2\pi}} e^{-\frac{x_{0.5}^2}{2\sigma^2}}. \tag{S8}$$

By cancelling the common amplitude constants and applying a natural logarithm, we arrive at

$$\ln\frac{1}{2} = -\ln 2 = -\frac{x_{0.5}^2}{2\sigma^2}. \tag{S9}$$

Rearranging and solving for $x_{0.5}$ we see that

$$x_{0.5} = \pm\sigma\sqrt{2\ln 2} \approx \pm 1.177\sigma \tag{S10}$$

and therefore the minimum distance between two objects which can be resolved is $\approx 2.355\sigma$. We can set this equal to the FWHM resolution limit to get

$$2\sigma\sqrt{\ln 4} = 0.51\frac{\lambda}{\text{NA}}. \tag{S11}$$

We can solve for $\sigma$ and get

$$\sigma = \frac{0.51\lambda}{2\sqrt{\ln 4}\text{NA}} \approx 0.217\frac{\lambda}{\text{NA}}, \tag{S12}$$

which, for the optimum GFP emission detection at 510nm, we arrive at

$$\sigma \approx \frac{110}{\text{NA}}\text{nm}, \tag{S13}$$

highlighting that NA and resolution are inversely proportional. From here, we have an equation which can directly influence the size of the Gaussian blur that we apply to account for the effects of microscopy, such that we can test various NA values of different lenses. For our microscopy assay, we have a lens with NA of 1.35, giving a blur radius of 82nm, around 90% of the CCD pixel size.
